# An evolutionary divergent thermodynamic brake in ZAP-70 fine-tunes the kinetic proofreading in T cell

**DOI:** 10.1101/2021.11.09.467998

**Authors:** Kaustav Gangopadhyay, Arnab Roy, Athira C. Chandradasan, Swarnendu Roy, Olivia Debnath, Soumee SenGupta, Subhankar Chowdhury, Dipjyoti Das, Rahul Das

## Abstract

T cell signaling starts with assembling several tyrosine kinases and adaptor proteins to the T cell receptor (TCR), following the antigen binding. The lifetime of the TCR: antigen complex and the time delay between the recruitment and activation of each kinase determines the T cell response. The mechanism by which the time delays are implemented in TCR signaling is not fully understood. Combining experiments and kinetic modeling, we here report a thermodynamic-brake in the regulatory module of ZAP-70, which determines the ligand selectivity, and may delay the ZAP-70 activation in TCR. Phylogenetic analysis revealed that the evolution of the thermodynamic-brake coincides with the divergence of the adaptive immune system to the cell-mediated and humoral responses. Paralogous kinase Syk expressed in B cells, does not possess such a functional thermodynamic brake, which may explain higher basal activation and lack of ligand selectivity by Syk.

## Introduction

The activation and quiescence in the cell-mediated immune response by T cell is regulated by a kinetic proofreading mechanism^1–3^. According to this mechanism, a time delay separates the binding of an antigen to the T cell receptor (TCR) from the subsequent downstream signaling^4–6^. TCR lacks intrinsic catalytic activity, and the downstream signaling starts with the recruitment of multiple kinases and adaptor proteins to the complex (Figure S1a) ^7–9^. Each recruitment step introduces a delay between the ligand binding and activation of the enzyme. The nonspecific interaction between self-antigen and TCR is short-lived and does not signal because the antigen: TCR complex dismantles before the activation of the downstream kinases. The interactions arising from foreign antigens are long-lived and get enough time to signal by activating the kinases. Paralogous kinases mediate the early events in B cell receptor (BCR) signaling of the humoral immune response employ conceptually similar mechanisms^10^. Nevertheless, the mechanism of differential activation of early T cell signaling compared to B cell remains unclear^11,12^.

The Syk family of non-receptor tyrosine kinases, ZAP-70 and Syk, are indispensable in the early stage of TCR and BCR signaling, respectively ^10,13^. Both the kinases are activated by recruiting to the membrane following antigen binding (Figure S1a). The dwell time of the kinases at the membrane determines their response^14^. ZAP-70 and Syk, both shares a modular structure composed of an N-terminal regulatory module and a C-terminal kinase domain (Figure 1a)^10,13^. The regulatory module is made up of tandem Src homology 2 (tSH2) domains connected by a helical linker called interdomain A (Figure 1a, c). In the inactive state (apo-state), the two SH2 domains adopt an ‘L’ like open conformation making them incompatible to ligand binding (Figure 1c)^15,16^. The Syk kinases are activated by binding to the doubly-phosphorylated immunoreceptor tyrosine-based activation (ITAM-Y2P) motifs at the TCR or BCR, respectively, through the tSH2 domain (Figure 1b, S1a)^17,18^. The tSH2 domain adopts a closed conformation upon binding to ITAM-Y2P, releasing the autoinhibitory interactions leading to the activation of the kinase domain (Figure 1c, S1a)^19,20^. The active conformation of ZAP-70 is stabilised by phosphorylating two key residues Y315 and Y319, by Lck, a Src family kinase, recruited to the antigen: TCR complex^21–23^. Several lines of evidence suggest that the ZAP-70 and Syk behave differently in the cell. Notably, ZAP-70 does not display basal activation, whereas Syk mediated basal signaling is essential for cell survival^24–29^. A ligand-independent closed conformation of the Syk tSH2 domain is proposed to facilitate high basal activation^30,31^. Despite the high sequence homology with Syk, it is not understood why the tSH2 domain of ZAP-70 could not adopt a stable closed conformation in the *apo*-state. Moreover, ZAP-70 displays a delayed Ca^2+^ response upon activation compared to Syk^12^. The tSH2 domain of ZAP-70 binds in a biphasic manner with a high degree of selectivity to a conserved ITAM-Y2P sequence, compared to hyperbolic binding in Syk^32–35^. The functional significance of biphasic ligand binding for T cell signaling remains unclear.

We present a kinetic model from a comparative study of the tSH2 domain of ZAP-70 and Syk, that explains the differential ligand-binding. We observed that the tSH2 domain of ZAP-70 binds to ITAM-Y2P in two-step kinetics, fast and slow, compared to one-step binding in Syk. The slow binding to the ZAP-70 tSH2 domain arises from a thermodynamic penalty (brake) that determines the ligand-selectivity and biases the conformational equilibrium of the *apo*-tSH2 domain towards the open conformation. Conversely, such thermodynamic break is non-functional in Syk tSH2-domain. The emergence of the thermodynamic brake coincides with the evolution of the BCR-TCR-MHC like immune system at the divergence of jawless and jawed fish approximately 500 million years ago^36^.

**Figure 1.**
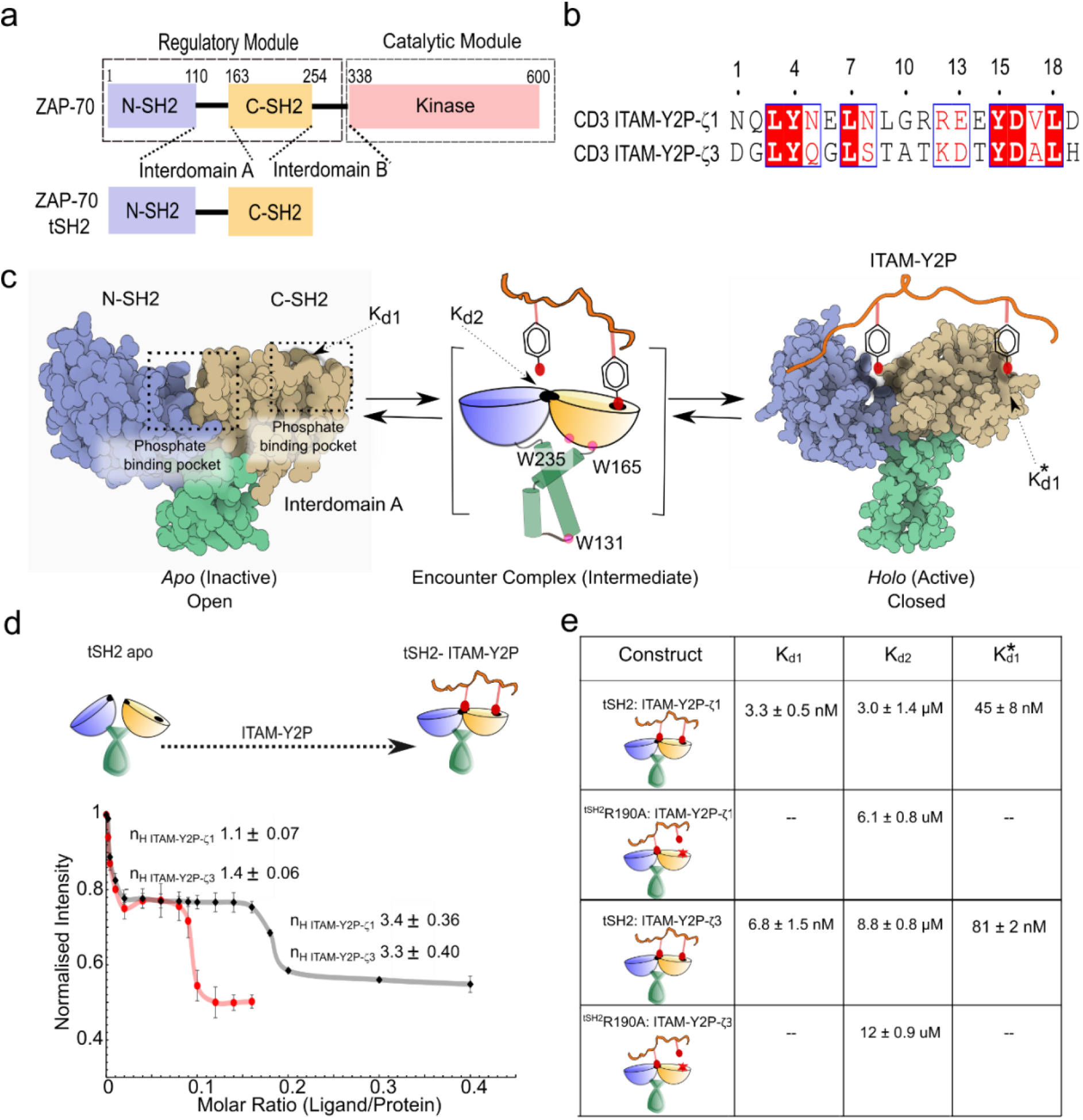
Binding of tSH2 domain of ZAP-70 to the ITAM-Y2P peptides: (a) Domain architecture of ZAP-70 full-length and the regulatory module. (b) Sequence alignment between ITAM-Y2P-ζ1 and ITAM-Y2P-ζ3 peptides. (c) Space-filled representation of tSH2-*apo* (PDB ID: 1M61) and tSH2-*holo* (PDB ID: 2OQ1) structure, the intermediate step is represented as cartoon. The N- and C-terminal SH2 domain, phosphate-binding pocket, and the respective binding constants are labeled. (d) Plot of change in intrinsic tryptophan fluorescence as a function of indicated ITAM-Y2P: tSH2 domain ratio. The Hill coefficients were calculated using Hill Plot. (e) Table summarizing the respective binding constants for the indicated tSH2 domain and ITAM-Y2P measured by ITC and fluorescence spectroscopy. Also see Figure S1.

## Results and Discussion

### The tSH2 domain of ZAP-70 is sensitive to the subtle changes in the ITAM peptide sequence

The ZAP-70 tSH2 domain binding to the doubly-phosphorylated ITAM-ζ1 peptide (ITAM-Y2P-ζ1) produce a biphasic curve with three distinct dissociation constants, *K*_*d*1_, *K*_*d*2_, and 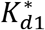 (Figure 1b–d) ^35^. First, the N-terminal phosphotyrosine residue from ITAM-Y2P binds uncooperatively to the C-SH2 phosphate-binding pocket (PBP) with a low nanomolar affinity (*K*_*d*1_) to form an encounter complex (Figure 1c–e). The formation of the tSH2: ITAM-Y2P encounter complex allows the assembly of the N-SH2 PBP. Subsequently, C-terminal phosphotyrosine residue from ITAM-Y2P binds weakly to the newly formed PBP with micromolar affinity (*K*_*d*2_). In the steady-state, the two binding events are interlinked by a plateau (Figure 1d). The second binding event remodels the C-SH2 PBP to an intermediate binding pocket 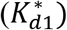 producing a *hill-coefficient* of 3.4 ± 0.36.

It was reported previously, the tSH2-domain of ZAP-70 displays hierarchical preference in binding to different ITAM sequences^17,33,37–39^. We begin by asking which part of the biphasic binding isotherm, in the steady-state, is sensitive to the ITAM peptide sequence (Figure 1b). We comparatively studied the binding of ITAM-Y2P-ζ3 to the tSH2 domain by intrinsic tryptophan fluorescence spectroscopy and isothermal titration calorimetry (ITC) (Figures 1b, d-e, S1b-f). We probed *K*_*d*1_, and 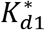 by fluorescence spectroscopy, and *K*_*d*2_ and 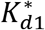 by ITC. To probe the *K*_*d*2_, we used ^tSH2^R190A mutant that impairs phosphotyrosine binding to C-SH2 PBP (Figures 1e, S1f).

We overserved that ITAM-Y2P-ζ3 binds weakly to the ZAP-70 tSH2 domain, compared to ITAM-Y2P-ζ1, consistent with the previous reports^17,40^ (Figure 1e). Our data revealed that the C-SH2 domain does not distinguish between the two ITAM-Y2P while forming the encounter complex. Both the peptides, ITAM-Y2P-ζ1 and ITAM-Y2P-ζ3, bind to the C-SH2 domain with low nanomolar affinity (*K*_*d*1_) of 3.3 ± 0.5 nM and 6.8 ± 1.5 nM, respectively. However, we noted a significant increase in the plateau width (Figure 1d). The ITAM-Y2P-ζ3 binding perturbed the *K*_*d*2_, and 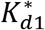 contributing to the overall increase in the dissociation constant (Figure 1e). We ask why the plateau-width in the steady-state binding (Figure 1d) is sensitive to the subtle changes in ligand type?

### A multistep ligand-receptor model explains the binding kinetics of ITAM-Y2P to ZAP-70

We developed a multistep mathematical-kinetic model to explain the biphasic bindings of ITAM-Y2P and tSH2 domains. This model comprised of different tSH2 domain conformations, open and closed, connected by a complex network (Figure 2a). The tSH2-*apo* state (open) ultimately reaches the tSH2-*holo* state (closed) through different pathways associated with distinct rates. The receptor adopts an *apo*-state, partially bound state (either C-SH2 or N-SH2 PBP are occupied), or *holo*-state, depending on the ligand concentration or type. A single binding event of one phosphotyrosine residue to either of the two PBP was modeled as first-order kinetics, meaning, the corresponding forward binding rate is proportional to the ligand concentration. We assumed, the transition between the open and closed conformations is ligand-independent (Figure 2a), and the formation of the encounter complex is the fastest (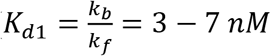; Figure 1e). To explain the biphasic binding (Figure 1d), we further assumed that the transitions to the *holo*-state from the partially bound states exhibit negative cooperativity. These steps represent kinetic penalties^41^, and occur with much slower rates (*w*_1_*k*_*f*_ and *w*_2_*k*_*f*_, with 0 < *w*_*i*_ < 1, *i* = 1 or 2) with dissociation constants 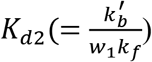 and 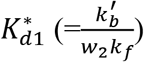, respectively. We anticipate, the variation in *K*_*d*1_, *K*_*d*2_, or 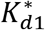 determine the steady-state response.

**Figure 2.**
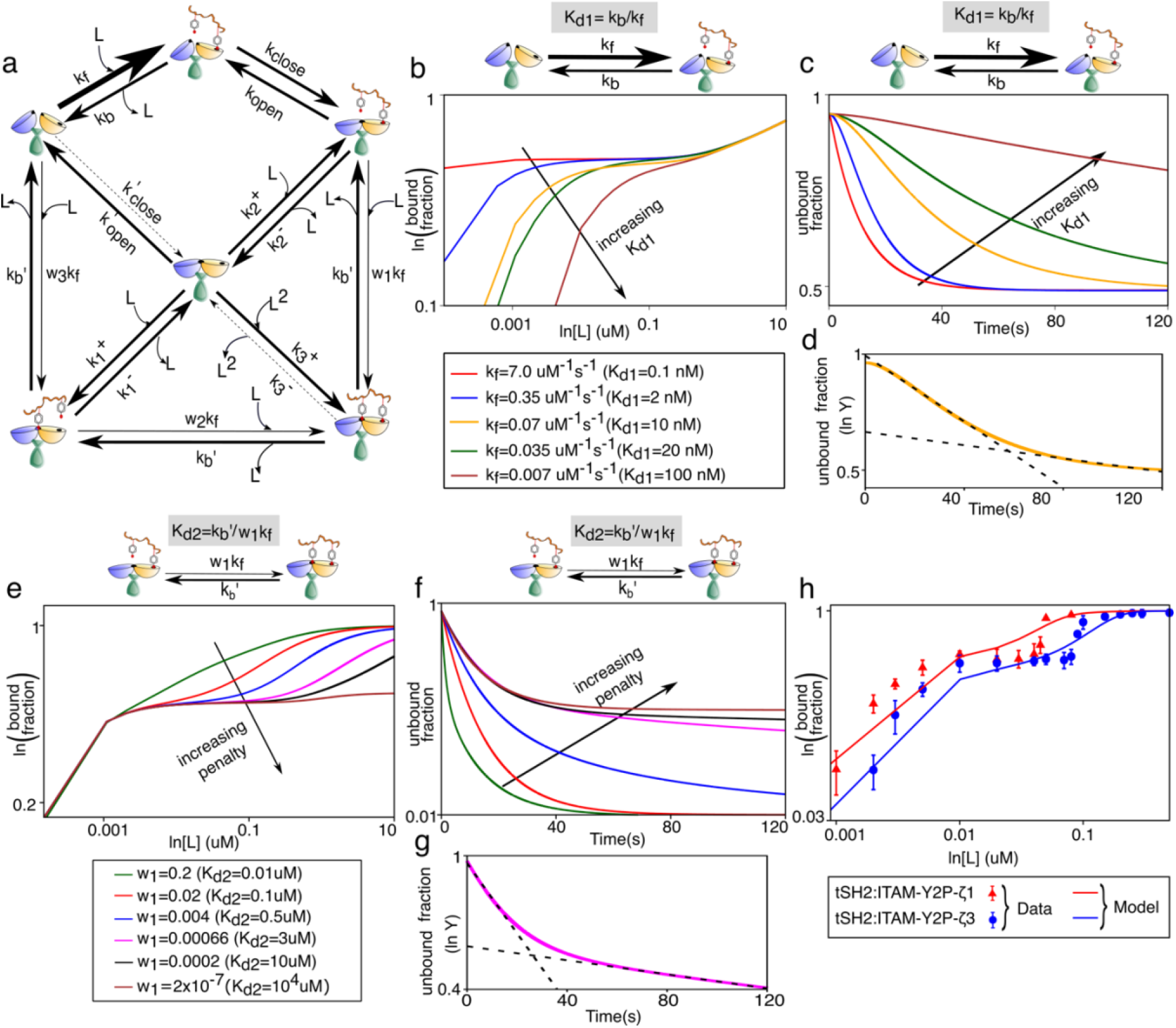
A kinetic-mathematical model explaining the biphasic ligand binding. **(a)** Schematic diagram of the model showing different reaction pathways. Arrow-widths represent distinct weightages of kinetic rates (bold arrows: higher rates, thin arrows: lower rates, and dotted arrows: negligible rates). **(b-c)** Effects of *K*_*d*1_ variation on the bound and unbound fraction under steady-state and pre-steady state, respectively. **(d)** Two-step decay is shown with two distinct exponential fits (dashed line) in a semi-log plot. (e-f) Effects of *K*_*d*2_ variation on the bound and unbound fraction under steady-state and pre-steady state, respectively. (g) Similar to panel d, two exponential fits (dashed line) represent distinctive two-step kinetics. (h) Comparison of theoretical prediction with the experimental data for binding of indicated ITAM-Y2P to the ZAP-70 tSH2 domains. Fitted parameters are in Table S5. Also see Figure S2.

We derived Ordinary Differential Equations (ODEs) for concentrations of chemical species using the law of mass-action (see Eq. S1). The ODEs were solved numerically to calculate the ligand bound fraction (defined in Eq. S2), and to predict the steady-state response and the kinetic behavior. In the steady-state, the *K*_*d*1_ determines the receptor sensitivity (initial rising slopes of the bound fractions) in low ligand concentration regime (nM level) and also modulates the plateau width (Figure 2b). The kinetic behavior mostly showed a single exponential decay except in a narrow intermediate range of *K*_*d*1_ (around 8nM – 20nM), where a two-step decay was observed (Figure 2c and 2d).

Next, we assessed the effect of *K*_*d*2_ (Figure 1c) variation by changing the penalty factor, *w*_1_ (Figure 2a). The *K*_*d*2_ variation modulated the ligand selectivity by altering the plateau width in the steady-state (Figure 2e). With a higher penalty (equivalently lower *w*_1_ or higher *K*_*d*2_), the plateau width became broader and displayed two-step kinetics with a sharp initial decay and a slow subsequent decrease of the unbound fraction. (Figure 2f–g).

The variation of 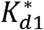 marginally alters the plateau width in the steady-state (Figure S2). Therefore, the slowest binding step to the N-SH2 PBP (*K*_*d*2_) mainly controls the plateau width. This conclusion correlates with the observed *K*_*d*2_, which is orders of magnitude lower than the *K*_*d*1_ and 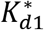 (Figure 1e), suggesting the corresponding step may impart a significant penalty in the dynamic binding of the ligand. However, measurement of free energy change is necessary to determine if a thermodynamic cost manifests as a kinetic penalty, as elucidated in the model.

Finally, for a particular set of parameter choices, the model prediction reasonably agreed with the experimental data of ITAM-Y2P-ζ1 and ITAM-Y2P-ζ3 bindings to the tSH2 domain (Figure 2h). In the model, reported values of *K*_*d*1_, *K*_*d*2_, and 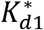 (Figure 1e) were used for the quantitative matching, but other parameters were unknown and chosen arbitrarily to fit the data (Table S5).

### The tSH2 domain of ZAP-70 binds ITAM-Y2P in two kinetic steps

We next probed the binding kinetics of ITAM-Y2P-ζ1 or ITAM-Y2P-ζ3 to the tSH2 domain by stop-flow fluorescence spectroscopy. We started with mixing excess ITAM-Y2P-ζ1 to the tSH2 domains at 10℃ and measured the change in tryptophan fluorescence intensity for 200 seconds (Figure 3a). We observed that the fluorescence intensity decay in two steps, fast (<200 ms) and slow (>20 s) (Figure 3a). Hence, we recorded all the kinetic experiments at two-time scales. All kinetic data were first normalized against the highest intensity and then subtracted by the blank (protein only sample) (Figure 3a–b) and fitted to a one-site association kinetics (Table 1).

**Table 1:**
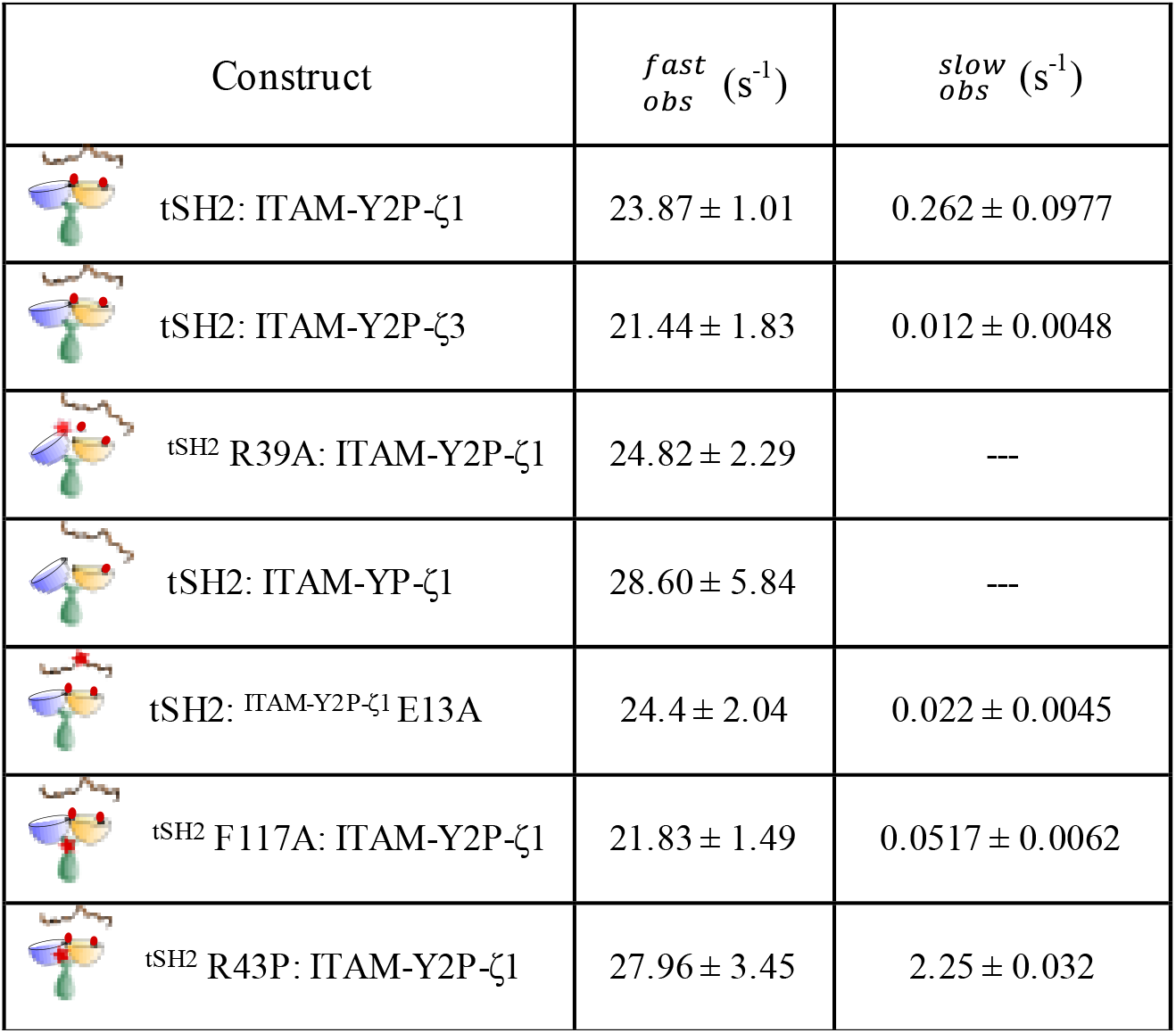
Observed rate for the ZAP-70 tSH2 domain and ITAM-Y2P binding

| Construct | $f_{obs}^{fast} (s^{-1})$ | $f_{obs}^{slow} (s^{-1})$ |
| --- | --- | --- |
| tSH2: ITAM-Y2P-ζ1 | $23.87 \pm 1.01$ | $0.262 \pm 0.0977$ |
| tSH2: ITAM-Y2P-ζ3 | $21.44 \pm 1.83$ | $0.012 \pm 0.0048$ |
| tSH2 R39A: ITAM-Y2P-ζ1 | $24.82 \pm 2.29$ | --- |
| tSH2: ITAM-YP-ζ1 | $28.60 \pm 5.84$ | --- |
| tSH2: ITAM-Y2P-ζ1 E13A | $24.4 \pm 2.04$ | $0.022 \pm 0.0045$ |
| tSH2 F117A: ITAM-Y2P-ζ1 | $21.83 \pm 1.49$ | $0.0517 \pm 0.0062$ |
| tSH2 R43P: ITAM-Y2P-ζ1 | $27.96 \pm 3.45$ | $2.25 \pm 0.032$ |

**Figure 3.**
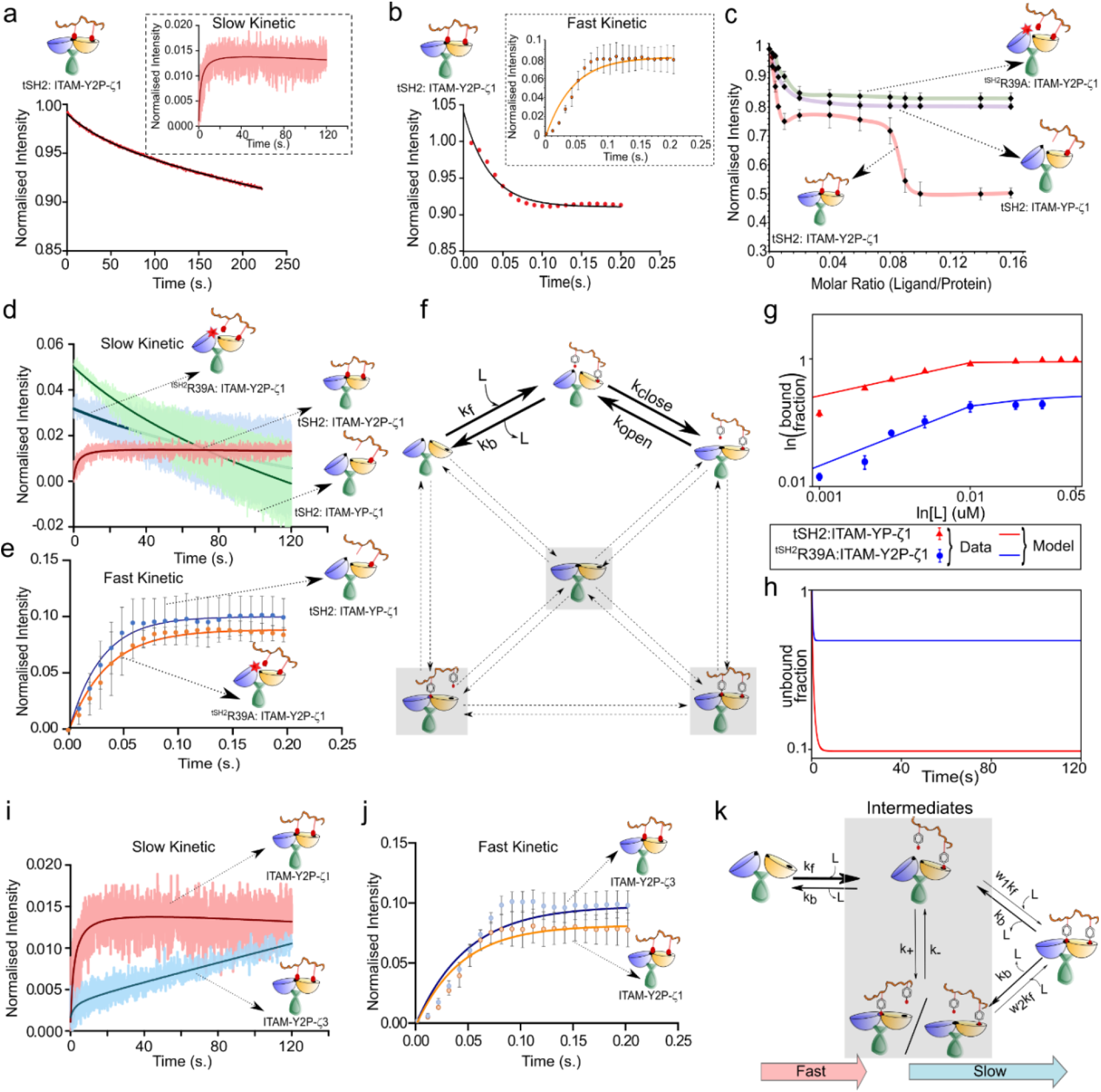
Binding kinetic of tSH2 domain of ZAP-70 and ITAM-Y2P. (a) and (b) the plot of time-dependent binding kinetics of tSH2 domain and 30μM ITAM-Y2P-ζ1 in the slow and fast time scales, respectively. Inset represents the single exponential fitting of the blank subtracted data. The error bar represents the standard deviation from three independent experiments. (c) The steady-state binding of the indicated tSH2 construct to ITAM-Y2P-ζ1 and ITAM-YP-ζ1. (d) and (e) The plots of slow and fast binding kinetics of indicated tSH2 construct to ITAM-Y2P-ζ1 and ITAM-YP-ζ1, respectively. (f) A modified version of the full model (Figure 2a). Forbidden intermediate conformations are highlighted by shadow boxes and dotted arrows. (g-h) The bound and unbound fractions of tSH2:ITAM-YP and ^tSH2^R39A:ITAM-Y2P under the steady state and pre-steady state conditions, respectively. In panel g, experimental steady-state data are also shown for comparison. (i-j) Binding kinetics of ITAM-Y2P-ζ1 and ITAM-Y2P-ζ3 to the ZAP-70 tSH2 domain. (k) A reduced kinetic model of ITAM binding. Also see Figure S3.

We observed that the ITAM-Y2P-ζ1 binds to the tSH2 domain with two observed rates of 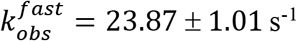, and 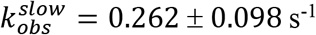 (Figure 3a–b, Table 1). Our mathematical model suggests that the fast-binding kinetic may arise (Figure 2d, 2g) during the formation of the encounter complex (*K*_*d*1_). To test that, we turn to two samples, one ^tSH2^R39A mutant that has an impaired N-SH2 PBP and second, a single phosphotyrosine ITAM-ζ1 peptide (ITAM-YP-ζ1) that will show only one binding event (Figure 3c). As expected, the steady-state fluorescence titration of ^tSH2^R39A to ITAM-Y2P-ζ1 and tSH2 domain to ITAM-YP-ζ1 showed that the first binding step is preserved with (*K*_*d*1_) of 8 ± 1.05 nM and 3.7 ± 0.1nM, respectively, and no subsequent binding was observed (Figures 3c, S3a-b). In the kinetics experiment, both the samples showed a 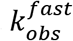 of 24.82 ± 2.29 s^−1^ and 28.60 ± 5.84 s^−1^, respectively, with no detectable slow binding (Figure 3d–e).

To check if our proposed model (Figure 2a) could explain the above data (Figure 3c–e), we introduced a modification in the model (Figure 3f). The N-SH2 binding is almost absent in both cases, and the partially bound state does not transform to the final *holo*-state. Hence, we ignore all pathways except the first encounter complex and the subsequent conformational change (Figure 3f). This assumption was sufficient for a quantitative matching between the model prediction and the experimental data in the steady-state (Figure 3g). The corresponding kinetic behaviour also showed a single exponential decay as observed in our experiment (Figure 3h).

Our model suggests that the tSH2 domain forms the encounter complex (*K*_*d*1_) with a fast-kinetic step, whereas the phosphate binding to the N-SH2 PBP (*K*_*d*2_) is the rate-limiting step and may determine the plateau width (Figure 2e–f). To validate, we measured the binding kinetics of the tSH2 domains and ITAM-Y2P-ζ3 (Figure 1d). Indeed, the ITAM-Y2P-ζ3 binds slower 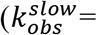 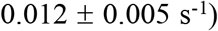 to the N-SH2 PBP compared to ITAM-Y2P-ζ1, with no significant change in the fast-kinetic step (Figure 2h, 3i–j, Table 1).

Since the *K*_*d*2_ seems to be the rate-limiting step, a reduced version of the full model (Figure 2a) is sufficient to capture essential features of the two-step kinetics. This reduced model has three basic steps: (i) a fast encounter to C-SH2 pocket (ii) followed by slower conformational changes of intermediates (open↔closed), and (iii) subsequent transition of intermediates to the final *holo*-state by imposing a kinetic penalty (Figure 3k). The minimal model was solved mathematically, and it produced qualitatively the same results as the full model (see Equations S4-S5, Figure S3 c-f). However, incorporating the finer details in the full model was necessary for quantitative matching with the data. We now ask, what determines the nature of the slow-kinetic step?

### The nature of the slow-kinetic step is determined by the structural coupling between the two SH2 domains

The ‘open to closed’ structural transitions of ZAP-70 tSH2 domain upon ligand binding requires cooperative interaction between the ITAM peptide, and the allosteric network resides in the tSH2 domain (Figure 1c)^17,35,42^. Analyzing the crystal structure of ZAP-70 tSH2 domain in complex with ITAM-Y2P-ζ1^19^, we identified a salt-bridge between the ^ITAM-ζ1^E13 and ^tSH2^K245 residue that may be critical for the structural coupling between the N- and C- SH2 domain during ligand binding (Figure 4a). The corresponding residue in ITAM-ζ3 is an aspartic acid, which may increase the distance between the ion-pair in the salt-bridge. This may reduce the transition rate to the final *holo*-state 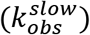, and increase the plateau width in the steady-state (Figure 1b, d, 3i). To test the role of the slat-bridge in determining the plateau width, we modulate the strength of the salt-bridge by changing the pH or by ^ITAM-ζ1^E13A mutation (Figure 4a–e, S4d, 1b). We observed that lowering the pH to 6.5 changes the surface potential of the tSH2 domain (Figure S4d) and marginally increases the plateau width, suggesting that the ^ITAM-ζ1^E13 and ^tSH2^K245 salt-bridge may be essential in the structural coupling (Figure 4b). To further investigate the role of the salt bridge, we measure the steady-state and kinetics of the tSH2 domain binding to the ^ITAM-ζ1^E13A. We observed that the ^ITAM-ζ1^E13A increases the plateau width in the steady-state binding with a significant increase in the *K*_*d*2_, and 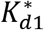 compared to the ITAM-Y2P-ζ1 (Figure 1e, 4c–f). The binding kinetics shows that the mutation in the ITAM-Y2P-ζ1 does not perturb the rate of encounter complex formation 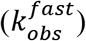, but slows down the transitioning to the closed-conformation 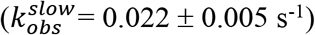, similar to ITAM-Y2P-ζ3 binding (Figure 3i–j, 4d–e, and Table 1). Our data suggest that the coupling between the two SH2 domains may determine the rate of the slow kinetic step 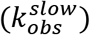.

**Figure 4.**
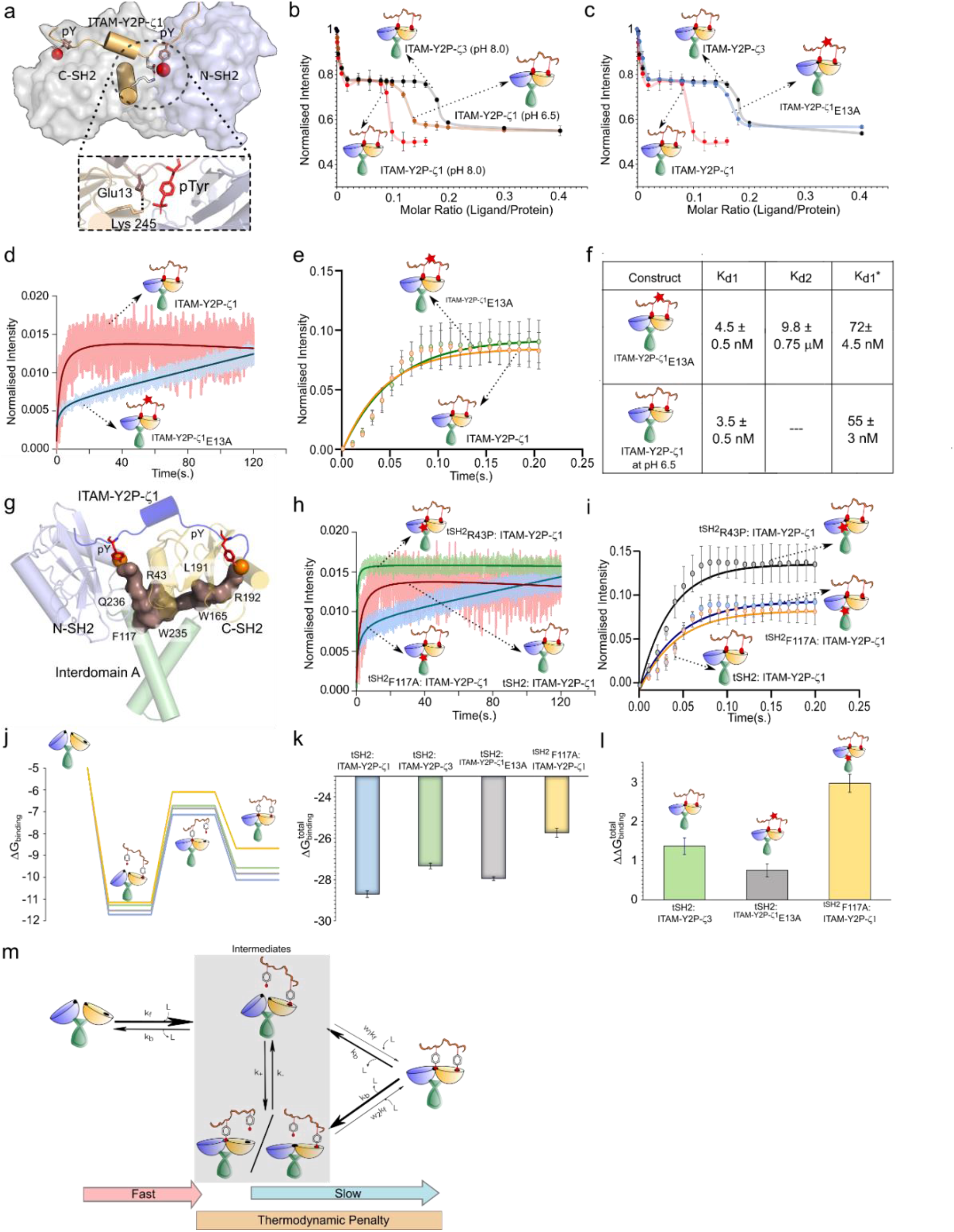
Structural evaluation of the tSH2 domain in determining the penalty step. (a) Space filled model of tSH2-*holo* structure of ZAP-70 (PDB ID: 2OQ1) highlighting the salt-bridge between the tSH2 domain and ITAM-Y2P-ζ1. (b) and (c) The steady-state binding of the tSH2 domain to ITAM-Y2P-ζ1 at indicated pH and ITAM-Y2P-ζ1 mutant, respectively. (d) and (e) The plots of slow and fast binding kinetics of tSH2 domain to indicated ITAM-Y2P-ζ1 peptides, respectively. (f) Table summarizing the dissociation constants for the indicated tSH2 domain and ITAM-Y2P samples. (g) Structure representing the allosteric network coupling the two SH2 domains in ZAP-70 (PDB ID: 2OQ1). (h) and (i) Slow and fast binding kinetics of indicated tSH2 domain mutants, respectively. The error bar represents the standard deviation from three independent experiments. (j) represents the change in Gibb’s free energy upon ligand binding (*ΔG*_*binding*_) in tSH2:ITAM-Y2P-ζ1 (blue), tSH2:ITAM-Y2P-ζ3 (green), tSH2:^ITAM-Y2P-ζ1^E13A (grey), and ^tSH2^F117A:ITAM-Y2P-ζ1 (yellow). (k) and (l) are the 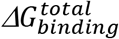 and 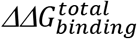, for the indicated tSH2 domain: ITAM-Y2P interactions. (m) Schematic model showing the binding kinetics and the thermodynamic penalty in the tSH2 and ITAM-Y2P interaction. Also see Figure S4.

A network of allosteric residues comprised of aromatic-aromatic stacking interaction allosterically couple the two SH2 domains of ZAP-70 during the transition to the final *holo*-conformation (Figure 4g).^35^ Mutating the residues in the allosteric network uncouples the formation of the encounter complex to the phosphotyrosine binding to the N-SH2 PBP (*K*_*d*2_). To determine if the mutation in the allosteric network will reduce the rate of conformational transition to the closed state 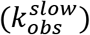, we studied the effect of ^tSH2^R43P and tSH2F117A mutants on the ITAM-Y2P-ζ1 binding kinetics (Figure 4g–i, table 1). We observed that the allosteric mutant did not perturb the rate of encounter complex formation 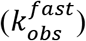 compared to the wild-type tSH2 domain. As anticipated, the allosteric mutants, ^tSH2^R43P, and ^tSH2^F117A, either inhibit or reduce the 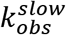, respectively (Figure 4h–i, Table 1). Analysis of the change in Gibb’s free-energy due to ligand binding (Δ*G*_*binding*_) suggests that the final transition to the close-conformation may impart a thermodynamic penalty (Figure 4j–l). Therefore, weakening the allosteric coupling led to a higher penalty at the transitions to the closed-state (Figure 3k). We conclude that the residues in the allosteric network collectively constitute a thermodynamic brake which may impart a delay between ZAP-70 binding and activation at the membrane (Figure 4m). It is widely believed that TCR shares a common ancestry and design principle with the B cell receptor (BCR). Is the thermodynamic brake present in Syk?

### Thermodynamic penalty coincides with the evolutionary-divergence of humoral and cell-mediated immune response

Syk is less selective than ZAP-70 and activated by a wide range of ITAM sequences in cells of innate and adaptive immune systems ^10,17,20,34,43–45^. Sequence analysis shows that most PBP and allosteric network residues in the regulatory module are conserved between Syk and ZAP-70 (Figure 5a, b). The exception is ^ZAP-70^R43; the corresponding residue in Syk is glutamine (Figure 5a, b). Phylogenetic mapping using the ZAP-70 as reference revealed that the ZAP-70 is conserved in all vertebrates and may appear first in the jawed fish (cartilage fish) (Figure 5c)^30,46^. Syk and Syk-related proteins are present in vertebrates and some invertebrates, including sponges and hydra (Figure 5c, S6a). This indicates, ZAP-70 may appear along with the evolution of the BCR-TCR-MHC like adaptive immune system at the divergence of jawless and jawed fish^36,47^. Intriguingly, the allosteric network residues comprising the thermodynamic brake in ZAP-70 are conserved, except residue L191 (Figure 5b–d). In amphibians and jawed fish, the Leu is often replaced by similar residues like Ile, Val, and Met. In Syk, two key residues in the allosteric network, N46, and W238 (corresponds to ZAP-70 R43 and W235, respectively), are not conserved (Figure 5b–d, S6b), suggesting that the thermodynamic brake may be non-functional.

**Figure 5:**
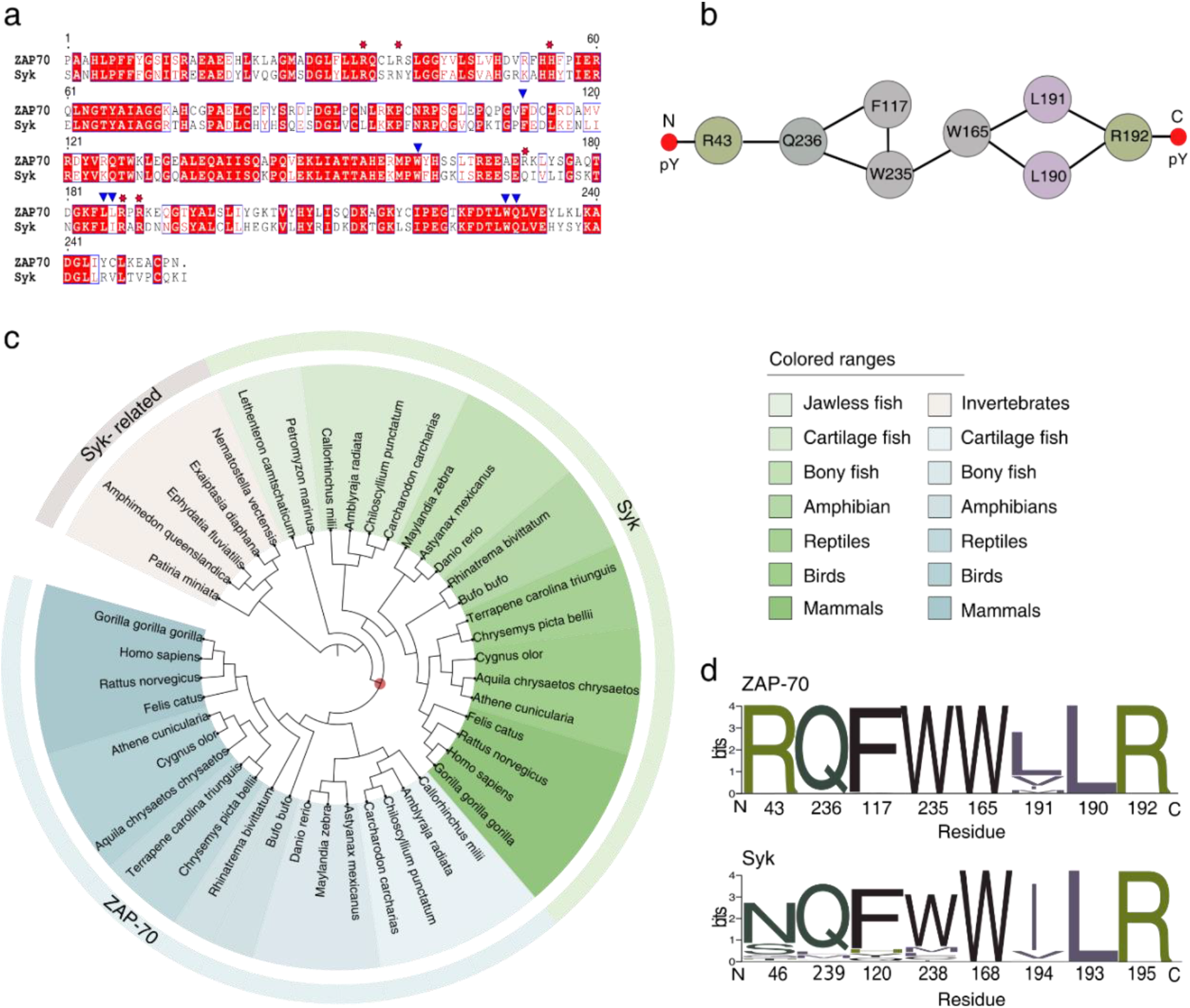
Phylogenetic analysis of allosteric network in the Syk family kinases. (a) Sequence Alignment of tSH2 domain of Syk and ZAP-70. The residues at the PBP are marked with red-star, and the allosteric network residues are marked with blue-arrow. (b) Schematic representation of undirected allosteric network in the tSH2 domain of ZAP-70. (c) Phylogenetic tree depicting the evolution of Syk family kinases, and Syk-related kinases. The emergence of ZAP-70 is marked as a red dot. (d) Sequence logo representing the conservation of allosteric network residue in ZAP-70 (top) and in Syk (including the Syk-related kinases) (bottom). Also see Table S1, Table S2, Figure S5 and S6.

To evaluate the role of the thermodynamic-brake in the Syk tSH2 domain, we characterized the steady-state binding of the tSH2 domain to ITAM-Y2P-ζ1, and ITAM-Y2P-ζ3, respectively. As reported previously, we observed that the Syk tSH2 domain could not distinguish between the ITAM sequences and bind with a similar *Kd* (~ 65 nM) (Figure 6a–b) 34. Unlike ZAP-70, the tSH2 domain of Syk binds ITAM-Y2P in a single fast kinetic step 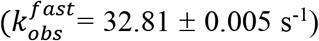, without any subsequent slow binding (Figure 6b–d). To evaluate if the absence of this slow binding correlates with the integrity of the allosteric network^35^, we inspected the open and closed structures of the Syk tSH2 domain (Figure 4g and 6e). Analysis of the crystal structures showed that the ZAP-70 tSH2 domain could adopt only two conformations, *open (apo)* and closed (holo) (Figure 1c)^19,31^. In comparison, the Syk tSH2 domain adopts three conformations: two structures in the *apo*-state, open and closed, and one ITAM-Y2P bound closed conformation^16,20,30^. In the apo-state of both Syk and ZAP-70, the SH2 domains are separated, preventing the aromatic amino acid residues to form the stacking interaction, which is central in coupling the N- and C- SH2 domains (Figure 4g, 6e). Surprisingly, the aromatic-aromatic stacking interaction does not form in the Syk tSH2 *holo*-state (Figure 6e). This indicates that the final transition to the *holo*-state does not require the allosteric network to assemble. To test that, we determine the steady-state binding of ITAM-Y2P-ζ1 to Syk tSH2 mutant, ^tSH2^F120A, and ^tSH2^W168C, respectively (Figure 5a, 6f–g). None of the mutants altered the ITAM-Y2P binding, implying that the allosteric network is non-functional in Syk.

**Figure 6.**
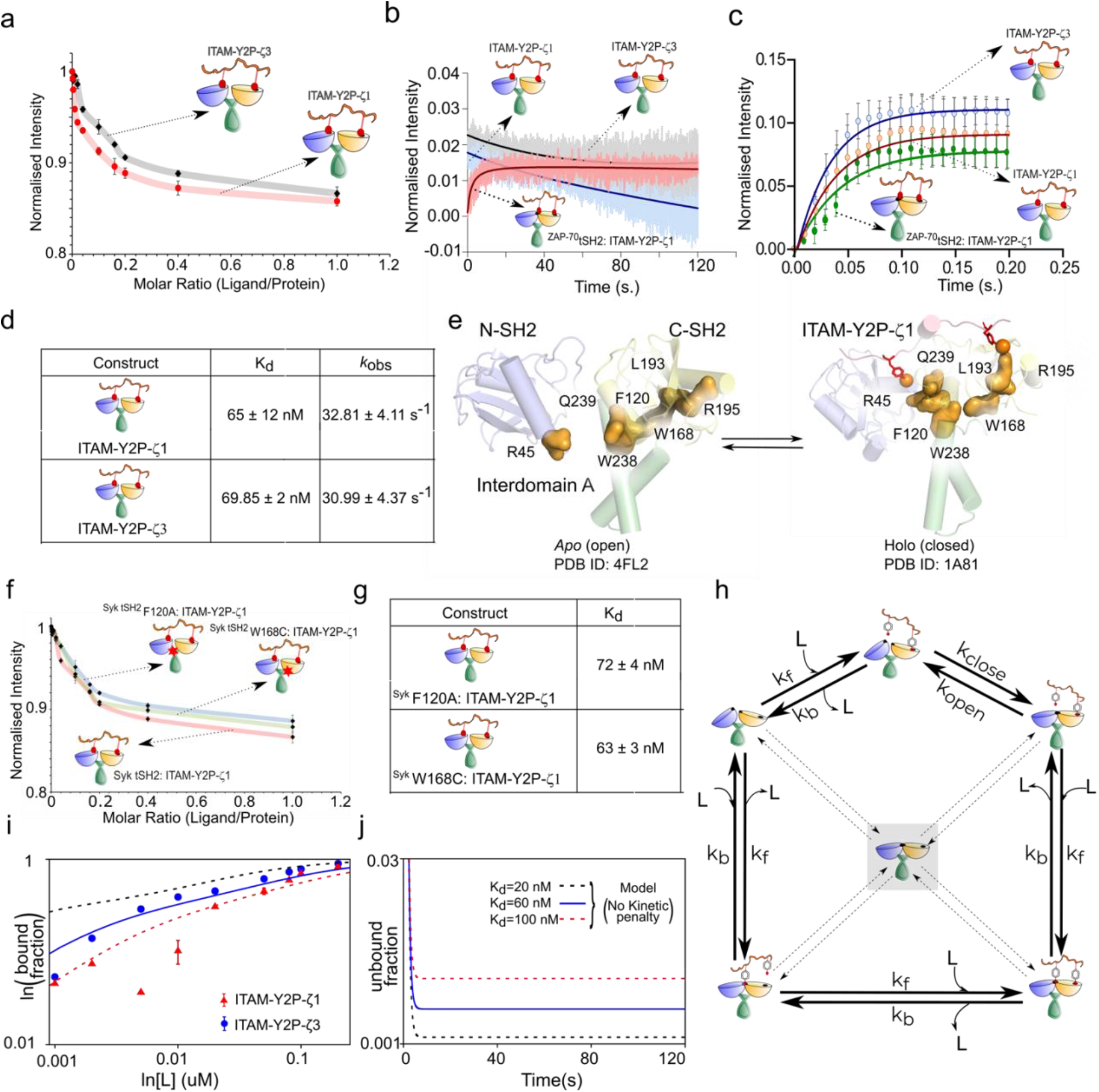
Steady-state and kinetic binding of Syk tSH2 domain to ITAM-Y2P: (a) Steady-state binding of ITAM-Y2P-ζ1 and ITAM-Y2P-ζ3 to the Syk tSH2 domain. (b) & (c) Slow and fast binding kinetics of indicated tSH2 domains to ITAM-Y2P peptides. (d) Table summarizing the dissociation constant (*K*_*d*_) and observed rate constant (*k*_*ods*_) for the Syk tSH2 domain. (e) Structure highlighting the allosteric network in the Syk tSH2 domain in *apo*-(PDB ID: 4FL2) and *holo-* state (PDB ID: 1A81), respectively. (f) Steady-state binding of indicated tSH2 construct to ITAM-Y2P-ζ1. (g) Tabulation of dissociation constant (*K*_*d*_) for the tSH2 allosteric mutant. The error bar represents the standard deviation from three independent experiments. (h) A modified version of the full model (Figure 2a) explaining Syk ITAM-Y2P interaction. Forbidden intermediate conformations are highlighted by shadow boxes and dotted arrows. (i) Comparison of the bound-fraction with the experimental data (filled circles and triangles) in the steady-state. (j) Pre-steady state kinetic profiles showing single-step decay. Also see Figure S7.

To explain if the absence of a penalty in Syk could lead to the hyperbolic binding in steady-state (Figure 6a) and one-step binding kinetics (Figure 6b–c), we remove the kinetic penalty in all pathways (*w*_*i*_ = 1) in the model described before (compare Figure 2a and 6h). We further assumed that the closed conformation in the *apo*-state (the central species in the network, Figure 6i) is short-lived. The experimental data of ITAM-Y2P binding to Syk matched with the modified model (Figure 6i), producing a single exponential decay (Compare Figure 6j and 2e). Therefore, the same kinetic model used for ZAP-70, in principle, also explains ligand binding in Syk if we consider independent binding in two SH2 domains^20,32^. This suggests that Syk tSH2 may have little or no penalty, explaining the lack of specificity for ITAM sequence^34,48^. Indeed, titration with ITAM-YP showed that Syk tSH2 domain binds with relatively stronger affinity (*K*_*d*_ = 27.7 ± 2.3 nM) compared to ITAM-Y2P (*K*_*d*_ = 65 ± 12 nM) (Figure 6a, S7a-b), suggesting the presence of negative cooperativity, as reported previously.^49^ We conclude, the thermodynamic brake is unique to ZAP-70, which may have coevolved in the adaptive immune cells during the divergence of the BCR-TCR-MHC like the immune system in the jawed fish.

## Conclusions

In summary, we have presented a unified kinetic model that explains the ZAP-70 and Syk tSH2 domain recruitment to the ITAM motifs at the membrane. Our model explains the recently observed closed conformation of an isolated *apo*-tSH2 domain structure of Syk^30^. Due to the absence of a penalty step, the Syk tSH2 domains could spontaneously adopt a closed conformation like in the *holo*-state, allowing the tSH2 domain to bind a wide range of ITAM-Y2P sequences with less selectivity. Absence of thermodynamic penalty facilitates binding of ITAM-Y2P even at lower concentrations observed under the basal level in B cells.

Our model suggests that incorporating a penalty step (thermodynamic brake) during the final transitioning to the *holo*-state (Figure 4m) is enough to explain the biphasic ITAM-Y2P binding and selectivity in the ZAP-70 tSH2 domain. The residues constituting the thermodynamic brake are conserved in all ZAP-70 kinases in the vertebrates, which coincides with the evolution of the humoral and cell-mediated immune system in jawed fish.

We propose that the thermodynamic brake is an essential component in the kinetic proofreading mechanism of ZAP-70 regulation. Several rate-limiting steps kinetically regulate the initiation of the ZAP-70-depended TCR signalling, including recruitment of coreceptor molecules coupled to Lck, dwell-time of ZAP-70, and phosphorylation of Y132 in LAT by ZAP-70 ^46,50,51^. In the T cell quiescence, the thermodynamic brake shifts the conformational equilibrium of the *apo*-tSH2 domain of ZAP-70 towards the open state. The additional energy barrier further stabilizes the compact inactive conformation of the kinase domain, which may shorten ZAP-70 dwell time at the TCR, causing reduced basal activation. The penalty step introduces a delay between the encounter complex formation and subsequent transition to the *holo*-tSH2 structure required for the activation of ZAP-70, which most likely explains delayed Ca^2+^ release in T cell^12^. The thermodynamic brake may provide an added layer of regulation in the kinetic proofreading, fundamentally differentiating TCR from BCR response.

## Materials and Methods

Details of materials and method comprising of the Purification of tSH2 domains, steady-state and pre-steady state Fluorescence experiments, Isothermal Titration Calorimetry, phylogenetic analysis, and Mathematical modeling can be found in Supplementary file.

## Supporting Information

The Supporting Information contain Material and Methods including Mathematical model, FigureS1-S7 and Table S1-S7.

## Author Contributions

The manuscript was written through the contribution of all authors. All authors have approved the final version of the manuscript. RD, and KG designed the experiments. KG, ACC, SR, OD, SSG and SC performed the Biochemical experiments and data analysis. ACC did the phylogenetic analysis of ZAP-70, OD, KG and RD performed the network analysis of tSH2 domain structure. DD, AR and ACC developed the mathematical model. RD, DD, KG and AR wrote the manuscript.

## Acknowledgements

The authors thank Prof. Pradipta Purkayastha for access to fluorimeter; Manas Pratim Chakraborty and Prosad Kumar Das for their valuable inputs. The authors thank Pritha Ganguly for her help with phylogenetic analysis. The authors thanks research funding from IISER Kolkata, infrastructural facilities supported by IISER Kolkata and DST-FIST (SR/FST/LS-II/2017/93(c)). This work is supported by grant from SERB (CRG/2020/000437) and DBT Ramalingaswami Fellowship (BT/RFF/Re-entry/14/2014) to RD; Ramalingaswami Fellowship (BT/RFF/Re-entry/52/2018) to DD. ACC and SSG are supported by the fellowships from CSIR-UGC, and UGC, respectively.

## Data Availability Statement

All the relevant data are contained within this article and in the supporting information.

## References

1. McKeithan, T.W. Kinetic proofreading in T-cell receptor signal transduction. Proceedings of the National Academy of Sciences 92, 5042–5046 (1995).

2. Rabinowitz, J.D., Beeson, C., Lyons, D.S., Davis, M.M. & McConnell, H.M. Kinetic discrimination in T-cell activation. Proceedings of the National Academy of Sciences 93, 1401–1405 (1996).

3. Chakraborty, A.K. & Weiss, A. Insights into the initiation of TCR signaling. Nature Immunology 15, 798–807 (2014).

4. Huang, W.Y.C. et al. Phosphotyrosine-mediated LAT assembly on membranes drives kinetic bifurcation in recruitment dynamics of the Ras activator SOS. Proceedings of the National Academy of Sciences 113, 8218–8223 (2016).

5. Tischer, D.K. & Weiner, O.D. Light-based tuning of ligand half-life supports kinetic proofreading model of T cell signaling. Elife 8(2019).

6. Yousefi, O.S. et al. Optogenetic control shows that kinetic proofreading regulates the activity of the T cell receptor. Elife 8(2019).

7. Gaud, G., Lesourne, R. & Love, P.E. Regulatory mechanisms in T cell receptor signalling. Nat Rev Immunol 18, 485–497 (2018).

8. Chan, A.C., Iwashima, M., Turck, C.W. & Weiss, A. ZAP-70: a 70 kd protein-tyrosine kinase that associates with the TCR zeta chain. Cell 71, 649–62 (1992).

9. Madrenas, J. et al. Zeta phosphorylation without ZAP-70 activation induced by TCR antagonists or partial agonists. Science 267, 515–8 (1995).

10. Mocsai, A., Ruland, J. & Tybulewicz, V.L. The SYK tyrosine kinase: a crucial player in diverse biological functions. Nat Rev Immunol 10, 387–402 (2010).

11. Borna, S., Fabisik, M., Ilievova, K., Dvoracek, T. & Brdicka, T. Mechanisms determining a differential threshold for sensing Src family kinase activity by B and T cell antigen receptors. J Biol Chem 295, 12935–12945 (2020).

12. Sadras, T. et al. Developmental partitioning of SYK and ZAP70 prevents autoimmunity and cancer. Mol Cell 81, 2094–2111.e9 (2021).

13. Au-Yeung, B.B., Shah, N.H., Shen, L. & Weiss, A. ZAP-70 in Signaling, Biology, and Disease. Annu Rev Immunol 36, 127–156 (2018).

14. Katz, Z.B., Novotna, L., Blount, A. & Lillemeier, B.F. A cycle of Zap70 kinase activation and release from the TCR amplifies and disperses antigenic stimuli. Nat Immunol 18, 86–95 (2017).

15. Deindl, S. et al. Structural basis for the inhibition of tyrosine kinase activity of ZAP-70. Cell 129, 735–46 (2007).

16. Gradler, U. et al. Structural and biophysical characterization of the Syk activation switch. J Mol Biol 425, 309–33 (2013).

17. Bu, J.Y., Shaw, A.S. & Chan, A.C. Analysis of the interaction of ZAP-70 and syk protein-tyrosine kinases with the T-cell antigen receptor by plasmon resonance. Proc Natl Acad Sci U S A 92, 5106–10 (1995).

18. Iwashima, M., Irving, B.A., van Oers, N.S., Chan, A.C. & Weiss, A. Sequential interactions of the TCR with two distinct cytoplasmic tyrosine kinases. Science 263, 1136–9 (1994).

19. Hatada, M.H. et al. Molecular basis for interaction of the protein tyrosine kinase ZAP-70 with the T-cell receptor. Nature 377, 32–8 (1995).

20. Futterer, K., Wong, J., Grucza, R.A., Chan, A.C. & Waksman, G. Structural basis for Syk tyrosine kinase ubiquity in signal transduction pathways revealed by the crystal structure of its regulatory SH2 domains bound to a dually phosphorylated ITAM peptide. J Mol Biol 281, 523–37 (1998).

21. Brdicka, T., Kadlecek, T.A., Roose, J.P., Pastuszak, A.W. & Weiss, A. Intramolecular regulatory switch in ZAP-70: analogy with receptor tyrosine kinases. Mol Cell Biol 25, 4924–33 (2005).

22. Yan, Q. et al. Structural basis for activation of ZAP-70 by phosphorylation of the SH2-kinase linker. Mol Cell Biol 33, 2188–201 (2013).

23. Williams, B.L. et al. Phosphorylation of Tyr319 in ZAP-70 is required for T-cell antigen receptor-dependent phospholipase C-gamma1 and Ras activation. Embo j 18, 1832–44 (1999).

24. Gururajan, M., Jennings, C.D. & Bondada, S. Cutting edge: constitutive B cell receptor signaling is critical for basal growth of B lymphoma. J Immunol 176, 5715–9 (2006).

25. Kraus, M., Alimzhanov, M.B., Rajewsky, N. & Rajewsky, K. Survival of Resting Mature B Lymphocytes Depends on BCR Signaling via the Igα/β Heterodimer. Cell 117, 787–800 (2004).

26. Chu, D.H. et al. The Syk protein tyrosine kinase can function independently of CD45 or Lck in T cell antigen receptor signaling. Embo j 15, 6251–61 (1996).

27. van Oers, N.S., Killeen, N. & Weiss, A. ZAP-70 is constitutively associated with tyrosine-phosphorylated TCR zeta in murine thymocytes and lymph node T cells. Immunity 1, 675–85 (1994).

28. Tsang, E. et al. Molecular mechanism of the Syk activation switch. J Biol Chem 283, 32650–9 (2008).

29. Goodfellow, H.S. et al. The catalytic activity of the kinase ZAP-70 mediates basal signaling and negative feedback of the T cell receptor pathway. Sci Signal 8, ra49 (2015).

30. Hobbs, H.T. et al. Differences in the dynamics of the tandem-SH2 modules of the Syk and ZAP-70 tyrosine kinases. Protein Science **n/a**.

31. Folmer, R.H., Geschwindner, S. & Xue, Y. Crystal structure and NMR studies of the apo SH2 domains of ZAP-70: two bikes rather than a tandem. Biochemistry 41, 14176–84 (2002).

32. Kumaran, S., Grucza, R.A. & Waksman, G. The tandem Src homology 2 domain of the Syk kinase: a molecular device that adapts to interphosphotyrosine distances. Proc Natl Acad Sci U S A 100, 14828–33 (2003).

33. Isakov, N. et al. ZAP-70 binding specificity to T cell receptor tyrosine-based activation motifs: the tandem SH2 domains of ZAP-70 bind distinct tyrosine-based activation motifs with varying affinity. J Exp Med 181, 375–80 (1995).

34. Grucza, R.A., Bradshaw, J.M., Mitaxov, V. & Waksman, G. Role of electrostatic interactions in SH2 domain recognition: salt-dependence of tyrosyl-phosphorylated peptide binding to the tandem SH2 domain of the Syk kinase and the single SH2 domain of the Src kinase. Biochemistry 39, 10072–81 (2000).

35. Gangopadhyay, K. et al. An allosteric hot spot in the tandem-SH2 domain of ZAP-70 regulates T-cell signaling. Biochemical Journal 477, 1287–1308 (2020).

36. Flajnik, M.F. & Kasahara, M. Origin and evolution of the adaptive immune system: genetic events and selective pressures. Nat Rev Genet 11, 47–59 (2010).

37. Labadia, M.E., Ingraham, R.H., Schembri-King, J., Morelock, M.M. & Jakes, S. Binding affinities of the SH2 domains of ZAP-70, p56lck and Shc to the zeta chain ITAMs of the T-cell receptor determined by surface plasmon resonance. J Leukoc Biol 59, 740–6 (1996).

38. Zenner, G., Vorherr, T., Mustelin, T. & Burn, P. Differential and multiple binding of signal transducing molecules to the ITAMs of the TCR-zeta chain. J Cell Biochem 63, 94–103 (1996).

39. Osman, N., Turner, H., Lucas, S., Reif, K. & Cantrell, D.A. The protein interactions of the immunoglobulin receptor family tyrosine-based activation motifs present in the T cell receptor zeta subunits and the CD3 gamma, delta and epsilon chains. Eur J Immunol 26, 1063–8 (1996).

40. Love, P.E. & Hayes, S.M. ITAM-mediated signaling by the T-cell antigen receptor. Cold Spring Harb Perspect Biol 2, a002485 (2010).

41. Sevlever, F., Di Bella, J.P. & Ventura, A.C. Discriminating between negative cooperativity and ligand binding to independent sites using pre-equilibrium properties of binding curves. PLOS Computational Biology 16, e1007929 (2020).

42. Clemens, L., Dushek, O. & Allard, J. Intrinsic Disorder in the T Cell Receptor Creates Cooperativity and Controls ZAP70 Binding. Biophysical Journal 120, 379–392 (2021).

43. Crowley, M.T. et al. A critical role for Syk in signal transduction and phagocytosis mediated by Fcgamma receptors on macrophages. J Exp Med 186, 1027–39 (1997).

44. Yanagi, S., Kurosaki, T. & Yamamura, H. The structure and function of nonreceptor tyrosine kinase p72syk expressed in hematopoietic cells. Cell Signal 7, 185–93 (1995).

45. Rogers, N.C. et al. Syk-dependent cytokine induction by Dectin-1 reveals a novel pattern recognition pathway for C type lectins. Immunity 22, 507–17 (2005).

46. Lo, W.L. et al. Slow phosphorylation of a tyrosine residue in LAT optimizes T cell ligand discrimination. Nat Immunol 20, 1481–1493 (2019).

47. Cooper, M.D. & Alder, M.N. The evolution of adaptive immune systems. Cell 124, 815–22 (2006).

48. Ottinger, E.A., Botfield, M.C. & Shoelson, S.E. Tandem SH2 domains confer high specificity in tyrosine kinase signaling. J Biol Chem 273, 729–35 (1998).

49. Feng, C. & Post, C.B. Insights into the allosteric regulation of Syk association with receptor ITAM, a multi-state equilibrium. Physical Chemistry Chemical Physics 18, 5807–5818 (2016).

50. Klammt, C. et al. T cell receptor dwell times control the kinase activity of Zap70. Nat Immunol 16, 961–9 (2015).

51. Stepanek, O. et al. Coreceptor scanning by the T cell receptor provides a mechanism for T cell tolerance. Cell 159, 333–45 (2014).

