## Supplementary information for "An evolutionary divergent thermodynamic brake in ZAP-70 fine-tunes the kinetic proofreading in T cell"

### MATERIALS AND METHOD:

#### Constructs:

ZAP-70 tSH2 wildtype (1-256), cloned in pSKB2 vector, was gifted from Prof. John Kuriyan, UC Berkeley. Syk tSH2 (7-263), cloned in pGEX6P1 vector, was gifted from Bruce Mayer (Addgene plasmid (Syk(NC)-SH2) #46521;). PCR based site-directed mutagenesis was done in the tSH2 background (R39A, R43P, F117A, W165C, R192A).

#### tSH2 domain expression and purification:

The tSH2 domain of ZAP-70 and Syk was expressed and purified as explained previously[1]. Briefly, the tSH2 domain of ZAP-70 was expressed in *E. coli* -BL21(DE3) cells by IPTG induction, and purified using Ni-NTA column. The eluted protein from the Ni-NTA column was further purified using a Q-column, followed by gel filtration chromatography. The purified tSH2 domain was buffer exchanged to 20 mM Tris-Cl, pH 8.0, 150 mM NaCl, 5 mM β-Mercaptoethanol, and 5% Glycerol and stored at -80˚C. The Syk tSH2 domain was expressed as Glutathione S-transferase (GST) fusion tag[2, 3] in *E. coli*-BL21(DE3) cells by IPTG induction. The protein was purified using Glutathione (GSH) column by eluting with reduced glutathione containing buffer (50 mM Tris-cl pH 8, 10 mM reduced glutathione, 10% Glycerol ). The GST tag was removed by digesting the protein with Prescission protease for overnight at 4°C. The digested sample was run over GSH column, followed by gel filtration chromatography for further purification. The purified constructs were stored in (20mM Tris-Cl, pH8.0, 150 mM NaCl, 5% Glycerol, and 5mM β-mercaptoethanol) at -80˚C.

#### Steady state fluorescence Experiments

Interaction between tSH2 domain and ITAM-Y2P in steady state were measured by following the change in intrinsic tryptophan fluorescence upon ITAM-Y2P binding by using PTI spectrofluorometer [1]. All the ITAM-Y2P peptides were purchased from Biotechdesk, and GMRF group labs. 0.5μM of tSH2 domain was titrated with various concentration of ITAM-Y2P peptide. The tryptophan fluorescence was recorded at $\lambda_{ex}$ 295nm and the emission was recorded from 300nm to 400nm. The dissociation constant ($K_d$) was derived by fitting the normalized intensity ($F_0$/F) versus ligand concentration using the following equation:

$$\frac{F_0}{F} = B_{max} * X^{n_H}/({K_d}^{n_H} + X^{n_H})$$

Where $F_0$ and F is fluorescence intensity in absence and presence of ligand, respectively. $B_{max}$ is the maximum binding, and nH is the Hill coefficient. The Hill coefficient (nH) was also determined from Hill plot[4]. The change in Gibb's free energy ($\Delta G_{binding}$) for each ligand binding step were calculated using $\Delta G_{binding}$ = -RTlnK, where K=1/$K_d$. The total change in Gibb's free energy $\Delta G_{binding}$ was calculated by adding the respective $\Delta G_{binding}$ at each step for respective tSH2 domain: ITAM-Y2P interaction. $\Delta\Delta G_{binding}$was calculated by subtracting $\Delta G_{binding}^{total}$ for the wildtype tSH2 domain: ITAM-Y2P-ζ1 from the indicated tSH2: ITAM-Y2P interactions (Figure 4l).

**Pre-steady state kinetics:**

The binding kinetics between tSH2 domain and ITAM-Y2P ligands were measured by following change in tryptophan fluorescence using a stop-flow fluorimeter (SFM2000bioLogic Spectrophotometer). The change in fluorescence intensity was measured at $\lambda_{ex}$ 290nm, $\lambda_{em}$ 350nm, and 10°C [5]. 100nM of protein (tSH2 domain) in 20 mM Tris (pH 8.0), 150 mM NaCl,

5% glycerol, 5 mM β-mercaptoethanol, and 30µM of ITAM-Y2P in same buffer, were mixed using syringe. Each transient was measured over 200s (for slow kinetics) or 1s (for fast kinetics), interval with 1500 and 101 time points, respectively. For each sample, a blank dataset was recorded by measuring the change in intensity for protein only. Each data set was normalized against the maximum intensity observed for the respective sample at time 0. The normalized sample was further corrected by blank subtraction. The observed rate ($k_{obs}$) was derived by fitting the change in intensity with respect to time by fitting to association kinetics equation implemented in Prism.

$$Y = Y_0 + (Y_{max} - Y_0)(1 - e^{-k20t})$$

Where, $Y_0$ is the intensity at time 0, $Y_{max}$ is the maximum intensity, $k$ is rate constant.

**Isothermal Titration calorimetry:**

Isothermal Titration Calorimetric studies were carried out using Malvern-PEAQ_ITC at 20°C. The wildtype tSH2 domain at a concentration of 20µM was titrated to different ITAM-Y2P constructs in glycerol free Tris buffer (20mM Tris-Cl, pH8.0, 150mM NaCl, 5mM β-mercaptoethanol). The $K_d$, ΔH, and ΔS were fitted into two-site sequential fit using ORIGIN as described previously [1]. In the ITC experiments with C-SH2 domain mutant (R190A), the protein was buffer exchanged to 20mM HEPES pH 8.2, 150mM NaCl nd 5mM β-mercaptoethanol. Titrations were carried out with 300µM of $^{\text{ITAM-Y2P-}\zeta 1}$EI3A, and ITAM-Y2P-ζ3 peptides and the data was fitted to one-site binding model to determine the $K_d$, ΔH and ΔS.

#### Phylogenetic analysis

The full-length protein sequences for Syk related kinases, Syk and ZAP-70 were acquired using multiple blast searches in the Uniprot database and PSI blast in NCBI. *Homo sapiens* Syk and ZAP-70 were separately used as query sequences against invertebrates and vertebrates. (Table S1 and Table S2)[6-8]. The secondary structures of proteins were verified using the PROSITE Expasy(Figure S5).[9] Only the proteins with tandem SH2 domains connected by a linker, which is connected to a kinase domain as seen in the *Homo sapiens* ZAP-70 with comparable interdomain A and B lengths were selected for sequence analysis. TK4, a Syk family kinase found in Platyhelminthes, was not considered for analysis due to considerably longer interdomain B (594 residues) than Syk kinase in *Homo sapiens* (110 residues). Also, HTK16 in *Hydra Vulgaris* and SHARK in *Drosophila* is not included in our analysis, since they possess Ankyrin repeats between the two SH2 domains. The protein tyrosine kinase in *Eptatretus burgeri* (jawless fish) was also removed as it comprises PH, Btk-type zinc finger, SH2, SH3, and Kinase domains. A total of 148 sequences from 82 organisms were selected, and accession numbers are summarized in Table S1 and S2. All the sequences were aligned, and a phylogenetic tree was constructed using MEGAX with the maximum likelihood method and bootstrapped 1000 times.[10-13] The phylogenetic tree was visualized using iTOL.[14] The WEB LOGO was used to visualize allosteric network residue conservation in Syk and ZAP-70.[15, 16].

### MATHEMATICAL ANALYSIS OF KINETIC MODELS:

Using laws of mass action, we derived the following set of ordinary differential equations (ODEs) from the kinetic model shown in Figure 2a (main text).

$$\left.\begin{aligned}
\frac{dR^{00}_{open}}{dt} &= k_b R^{01}_{open} + k_b^{'} R^{10} + k'_{open} R^{00}_{closed} - (1 + w_3) k_f L R^{00}_{open} - k'_{close} R^{00}_{open} \\
\frac{dR^{00}_{closed}}{dt} &= k'_{close} R^{00}_{open} + k_1^{-} R^{10} + k_2^{-} R^{01} + k_3^{-} R^{11} - k'_{open} R^{00}_{closed} - (k_1^{+} + k_2^{+}) L R^{00}_{closed} - k_3^{+} L^2 R^{00}_{closed} \\
\frac{dR^{01}_{open}}{dt} &= k_f L R^{00}_{open} + k_{open} R^{01} - (k_b + k_{close}) R^{01}_{open} \\
\frac{dR^{01}}{dt} &= k_{close} R^{01}_{open} + k_b^{'} R^{11} + k_2^{+} L R^{00}_{closed} - k_{open} R^{01} - k_2^{-} R^{01} - w_1 k_f L R^{01}
\end{aligned}\right\} \quad \text{(S1)}$$

$$\frac{dR^{10}}{dt} = w_3 k_f L R^{00}_{open} + k_b^{'} R^{11} + k_1^{+} L R^{00}_{closed} - k_b^{'} R^{10} - k_1^{-} R^{10} - w_2 k_f L R^{10}$$

$$\frac{dR^{11}}{dt} = w_1 k_f L R^{01} + w_2 k_f L R^{10} + k_3^{+} L^2 R^{00}_{closed} - 2 k_b^{'} R^{11} - k_3^{-} R^{11}$$

In the above set of equations, $R^{ij}_{open}$ and $R^{ij}_{closed}$ denote the concentrations of receptors in different conformations (see Table S3 for the meaning of symbols). Here i and j denote the binding (i,j=1) or unbinding (i,j=0) at the N-SH2 and C-SH2 binding sites respectively. Further, L denotes the concentration of the ligand. We assumed a single binding event at any of the two sites as 1st order kinetics (i.e., binding rate is proportional to L) and double binding events as 2nd order kinetics (i.e., proportional to $L^2$). We assumed the species $R^{00}$ and $R^{01}$ can be either in open or closed conformation, while $R^{10}$ and $R^{11}$ are always in closed conformation (See Figure 2a and Table S3).

Following Sevlever *et al.* (Plos Comp. Bio., 2020), we define the fraction of partially and fully bound species by the 'bound fraction' as below: (S2)

$$\Phi_b = \frac{\left(\frac{1}{2}R^{01} + \frac{1}{2}R^{10} + R^{11}\right)}{R_{total}}, \text{ where } R_{total} = \left(R^{00}_{closed} + R^{00}_{open} + R^{01}_{open} + R^{01} + R^{10} + R^{11}\right)$$

Conversely, the 'unbound fraction' is given by, $\Phi_{ub} = (1 - \Phi_b)$

We solved the above set of ODEs numerically in Mathematica (using Parametric NDSOLVE) and obtained steady state concentrations as the function of ligand concentrations at a large time. We assumed that the steady state has reached when all concentrations were constant over time (we took numerical data at t >> 5 hour). From steady state concentrations, we calculated the bound fraction in Figure 2b and 2e. The initial condition was: $R^{00}_{open}$ = 500 nM and $R^{00}_{closed} = R^{01}_{open} = R^{01} = R^{10} = R^{11} = 0$, at t=0.

To observe the kinetic behavior, we numerically solved the same set of ODEs (Eq. S1). We used $R^{00}_{open}$ = 500 nM and L = 10 uM, ($R^{00}_{closed} = R^{01}_{open} = R^{01} = R^{10} = R^{11} = 0$) initially for Figure 2f and 2g; while $R^{00}_{open}$ = 100 nM and L = 500 nM, ($R^{00}_{closed} = R^{01}_{open} = R^{01} = R^{10} = R^{11} = 0$) initially for Figure 2c and 2d. All other parameter values used in the model to produce Figure 2b-2h are summarized in Tables S4 and S5. Note, when $k_f$ was varied in Figure 2b-c, we kept the rates fixed in all penalty-inducing steps, i.e., we fixed the values of $w_i k_f$ by proportionately changing $w_i$ $(i = 1,2,3)$.

To match the theoretical bound fraction with the experimental data, we calculated the bound fraction from the experiment as below:

$$\Phi_b \text{ (experimental)} = \frac{(F_{max} - F)}{(F_{max} - F_{min})} \,, \tag{S3}$$

where F is the fluorescence intensity and $F_{max}$ and $F_{min}$ are maximum and minimum values of the intensity, respectively.

The steady-state behavior of tSH2: ITAM-YP-ζ1 and $^{tSH2}$R39A: ITAM-Y2P-ζ1 both were explained by the modified model (see Figure 3f). Mathematically, we solved the same set of ODEs (Eq. S1) at appropriate limits (see Table S6). In particular, we set $w_1 = w_2 = w_3 = 0$ and $\{k'_b, k_1^+, k_1^-, k_2^+, k_2^-, k_3^+, k_3^-, k'_{close}, k'_{open}\} \sim 0$ (i.e., negligible). In these cases, since only the C-SH2 binding site can be occupied, the bound fraction would be:

$$\Phi_b = \left(\frac{R^{01}}{R_{total}}\right)$$

We numerically calculated $\Phi_b$ and $\Phi_{ub}$ and plotted them in Figure 3g-3h. The corresponding parameter values used to match the theoretical bound fraction with the experimental data are mentioned in Table S6. According to the experimental data, C-SH2 binding affinity are similar for tSH2: ITAM-YP-ζ1 and $^{tSH2}$R39A: ITAM-Y2P-ζ1 ($K_{d1}$= 4 nM and 8 nM respectively; Figure S3b). However, we had to assume that the rate of closed to open conformational change for tSH2: ITAM-YP-ζ1 should be roughly 20 times faster than the $^{tSH2}$R39A: ITAM-Y2P-ζ1 for quantitative matching between the experimental and theoretical bound fraction. This suggests that the $^{tSH2}$R39A: ITAM-Y2P-ζ1 complex is relatively unstable than tSH2: ITAM-YP-ζ1.

Similar to the model in Figure 2a, we have derived the ODEs for the reduced model shown in Figure 3k. The ODEs are:

$$\left.\begin{aligned}
\frac{d\mathrm{R}^{00}_{\mathrm{open}}}{dt} &= k_b \mathrm{R}^{01}_{\mathrm{open}} - k_f \mathrm{L}\mathrm{R}^{00}_{\mathrm{open}} \\
\frac{d\mathrm{R}^{01}_{\mathrm{open}}}{dt} &= k_f \mathrm{L}\mathrm{R}^{00}_{\mathrm{open}} + k_- \mathrm{R}^{\mathrm{ij}}_{\mathrm{closed}} + k_b \mathrm{R}^{11} \text{-} (k_b + k_+ + w_1 k_f \mathrm{L})\mathrm{R}^{01}_{\mathrm{open}} \\
\frac{d\mathrm{R}^{\mathrm{ij}}_{\mathrm{closed}}}{dt} &= k_+ \mathrm{R}^{01}_{\mathrm{open}} + k_b \mathrm{R}^{11} - (k_- + w_2 k_f \mathrm{L})\mathrm{R}^{\mathrm{ij}}_{\mathrm{closed}}
\end{aligned}\right\} \quad \text{(S4)}$$

$$\frac{d\mathrm{R}^{11}}{dt} = w_1 k_f \mathrm{L}\mathrm{R}^{01}_{\mathrm{open}} + w_2 k_f \mathrm{L}\mathrm{R}^{\mathrm{ij}}_{\mathrm{closed}} - 2k_b \mathrm{R}^{11}$$

Here, $\mathrm{R}^{\mathrm{ij}}_{\mathrm{closed}}$ denote all possible intermediates which are in partially-bound close conformations. On the other hand, $\mathrm{R}^{11}$ is the final holo-state (closed). Similar to the full model, the switching between open to close conformations (via rates $k_+$ and $k_-$) are ligand independent.

Solving the above set of simple ODEs, we derived exact expressions for the steady state concentrations as follows:

$$\left.\begin{aligned}
\mathrm{R}^{01}_{\mathrm{open}} &= \left(\frac{\mathrm{L}}{K_{d1}}\right) \times \mathrm{R}^{00}_{\mathrm{open}} \\
\mathrm{R}^{\mathrm{ij}}_{\mathrm{closed}} &= \left(\frac{2k_+ + w_1 k_f \mathrm{L}}{2k_- + w_2 k_f \mathrm{L}}\right) \times \left(\frac{\mathrm{L}}{K_{d1}}\right) \times \mathrm{R}^{00}_{\mathrm{open}} \\
\mathrm{R}^{11} &= \left(\frac{w_1 k_- + w_2\left(k_+ + w_1 k_f \mathrm{L}\right)}{2k_- + w_2 k_f \mathrm{L}}\right) \times \left(\frac{\mathrm{L}}{K_{d1}}\right)^2 \times \mathrm{R}^{00}_{\mathrm{open}}
\end{aligned}\right\} \quad \text{(S5)}$$

where, $K_{d1} = \frac{k_b}{k_f}$

We used the same values for $k_f$, $k_b$, $w_1$, $w_2$ as in Table S4, while we took $k_+$ = 0.00007/s, $k_-$=0.0007/s ($\frac{k_-}{k_+} = 10$). Using these parameters, from the above expressions (Eq. S5) we calculated $\Phi_b$ and $\Phi_{ub}$ and plotted in Figure S3 c-f. These figures show qualitatively the same results as in the full model (compare Figure 2 b-f, and Figure S3 c-f), suggesting the full model and the reduced model are qualitatively similar.

Binding of ITAM-Y2P-ζ1 and ITAM-Y2P-ζ3 with Syk were also explained by modification in the same mathematical model (Figure 6h). We solved the same set of ODEs (Eq. S1) with the conditions $w_1 = w_2 = w_3 = 1$ (i.e., no kinetic penalty) and $\{k_1^+, k_1^-, k_2^+, k_2^-, k_3^+, k_3^-, k'_{close}, k'_{open}\}$[8] ~ 0 (i.e. negligible). We numerically calculated $\Phi_b$ and $\Phi_{ub}$ and showed in Figure 6i-6j. Parameters corresponding to these plots are shown in Table S7. According to the experiments, Syk cannot distinguish between ITAM-Y2P-ζ1 and ITAM-Y2P-ζ3 and show the same $K_d$ values (around 65-70nM). That's why we varied $K_d$ values from 20nM to 100nM range to have a quantitative match between the theoretical and the experimental bound fraction.

**Mathematica Codes:** all codes are publicly available online at the following link [https://github.com/arnabroy97/ODE_calculations](https://github.com/arnabroy97/ODE_calculations)

**Table S1:** Accession number of Syk and Syk Related Kinases of different organisms

| Sl. no | Accession number | Gene | Species | Phylum | Class | Domains |
|---|---|---|---|---|---|---|
| 1 | Q9Y1X9 | EfPTK151 | *Ephydatia fluviatilis* | Porifera | Demospongiae | 2 SH2, 1 kinase |
| 2 | A0A1X7V2S5 | 100640238 | *Amphimedon queenslandica* | Porifera | Demospongiae | 2 SH2, 1 kinase |
| 3 | O77440 | HTK98 | *Hydra vulgaris* | Cnidaria | Hydrozoa | 2 SH2, 1 kinase |
| 4 | A0A2B4SFM1 | SYK | *Stylophora pistillata* | Cnidaria | Anthozoa | 2 SH2, 1 Kinase |
| 5 | A0A6P8ILN3 | LOC116302664 | *Actinia tenebrosa* | Cnidaria | Anthozoa | 2 SH2, 1 Kinase |
| 6 | XP_020903367.1 | SYK IsoformX1 | *Exaiptasia diaphana* | Cnidaria | Anthozoa | 2 SH2, 1 Kinase |
| 7 | XP_032226357.1 | SYK IsoformX1 | *Nematostella vectensis* | Cnidaria | Anthozoa | 2 SH2, 1 Kinase |
| 8 | XP_028390911.1 | SYK IsoformX1 | *Dendronephthya gigantea* | Cnidaria | Anthozoa | 2 SH2, 1 Kinase |
| 9 | XP_021340359.1 | SYK-like | *Mizuhopecten yessoensis* | Mollusca | Bivalvia | 2 SH2, 1 Kinase |
| 10 | CAG2209972.1 | SYK | *Mytilus edulis* | Mollusca | Bivalvia | 2 SH2, 1 Kinase |
| 11 | XP_033736612.1 | SYK-like | *Pecten maximus* | Mollusca | Bivalvia | 2 SH2, 1 Kinase |
| 12 | XP_041360389.1 | SYK-like | *Gigantopelta aegis* | Mollusca | Gastropoda | 2 SH2, 1 Kinase |
| 13 | XP_038067744.1 | SYK IsoformX1 | *Patiria miniata* | Echinodermata | Asteroidea | 2 SH2, 1 Kinase |
| 14 | XP_030851825.1 | SYK IsoformX1 | *Strongylocentrotus purpuratus* | Echinodermata | Echinoidea | 2 SH2, 1 Kinase |
| 15 | XP_032816776.1 | SYK IsoformX1 | *Petromyzon marinus* | Chordata | Agnatha | 2 SH2, 1 Kinase |
| 16 | A0A0C5DLZ7 | SYK | *Lethenteron camtschaticum* | Chordata | Agnatha | 2 SH2, 1 Kinase |
| 17 | A0A4W3IP53 | SYK | *Callorhinchus milii* | Chordata | Chondrichthyes | 2 SH2, 1 Kinase |
| 18 | A0A401SZZ3 | chiPu_0014454 | *Chiloscyllium punctatum* | Chordata | Chondrichthyes | 2 SH2, 1 Kinase |
| 19 | XP_041042106.1 | SYK IsoformX1 | *Carcharodon carcharias* | Chordata | Chondrichthyes | 2 SH2, 1 Kinase |
| 20 | XP_032873635.1 | SYK IsoformX1 | *Amblyraja radiata* | Chordata | Chondrichthyes | 2 SH2, 1 Kinase |
| 21 | XP_038660658.1 | SYK IsoformX1 | *Scyliorhinus canicula* | Chordata | Chondrichthyes | 2 SH2, 1 Kinase |
| 22 | XP_008323672.1 | SYK IsoformX1 | *Cynoglossus semilaevis* | Chordata | Osteichthyes | 2 SH2, 1 Kinase |
| 23 | A0A672HYJ7 | SYK | *Salarias fasciatus* | Chordata | Osteichthyes | 2 SH2, 1 Kinase |
| 24 | XP_014191223.1 | SYK IsoformX1 | *Haplochromis burtoni* | Chordata | Osteichthyes | 2 SH2, 1 Kinase |
| 25 | XP_015191917.1 | SYK IsoformX1 | *Lepisosteus oculatus* | Chordata | Osteichthyes | 2 SH2, 1 Kinase |
| 26 | H3ATL1 | SYK | *Latimeria chalumnae* | Chordata | Osteichthyes | 2 SH2, 1 Kinase |
| 27 | XP_026152626.1 | SYK | *Mastacembelus armatus* | Chordata | Osteichthyes | 2 SH2, 1 Kinase |
| 28 | RXM97385.1 | SYK | *Acipenser ruthenus* | Chordata | Osteichthyes | 2 SH2, 1 Kinase |
| 29 | NP_998008.2 | SYK | *Danio rerio* | Chordata | Osteichthyes | 2 SH2, 1 Kinase |
| 30 | A0A672MPJ9 | SYK | *Sinocyclocheilus grahami* | Chordata | Osteichthyes | 2 SH2, 1 Kinase |
| 31 | KAG9263497.1 | SYK | *Astyanax mexicanus* | Chordata | Osteichthyes | 2 SH2, 1 Kinase |
| 32 | XP_026798305.1 | SYK | *Pangasianodon hypophthalmus* | Chordata | Osteichthyes | 2 SH2, 1 Kinase |
| 33 | XP_012711475.1 | SYK IsoformX1 | *Fundulus heteroclitus* | Chordata | Osteichthyes | 2 SH2, 1 Kinase |

| | | | | | | |
|---|---|---|---|---|---|---|
| 34 | XP_004564629.1 | SYK | *Maylandia zebra* | Chordata | Osteichthyes | 2 SH2, 1 Kinase |
| 35 | XP_018600403.1 | SYK IsoformX1 | *Scleropages formosus* | Chordata | Osteichthyes | 2 SH2, 1 Kinase |
| 36 | A0A1L8HYI7 | SYK.L | *Xenopus laevis* | Chordata | Amphibia | 2 SH2, 1 Kinase |
| 37 | F7DXJ8 | SYK | *Xenopus tropicalis* | Chordata | Amphibia | 2 SH2, 1 Kinase |
| 38 | A0A6P8PLK0 | SYK | *Geotrypetes seraphini* | Chordata | Amphibia | 2 SH2, 1 Kinase |
| 39 | A0A6P7X829 | SYK | *Microcaecilia unicolor* | Chordata | Amphibia | 2 SH2, 1 Kinase |
| 40 | XP_040272878.1 | SYK IsoformX1 | *Bufo bufo* | Chordata | Amphibia | 2 SH2, 1 Kinase |
| 41 | XP_029474555.1 | SYK IsoformX1 | *Rhinatrema bivittatum* | Chordata | Amphibia | 2 SH2, 1 Kinase |
| 42 | A0A6J0U1P1 | SYK | *Pogona vitticeps* | Chordata | Reptile | 2 SH2, 1 Kinase |
| 43 | XP_005307710.1 | SYK IsoformX1 | *Chrysemys picta bellii* | Chordata | Reptile | 2 SH2, 1 Kinase |
| 44 | A0A402FJ14 | parPi_0018852 | *Paroedura picta* | Chordata | Reptile | 2 SH2, 1 Kinase |
| 45 | XP_024070757.2 | SYK IsoformX1 | *Terrapene carolina triunguis* | Chordata | Reptile | 2 SH2, 1 Kinase |
| 46 | A0A670JJY3 | SYK | *Podarcis muralis* | Chordata | Reptile | 2 SH2, 1 Kinase |
| 47 | A0A6I9XWC1 | SYK | *Thamnophis sirtalis* | Chordata | Reptile | 2 SH2, 1 Kinase |
| 48 | A0A674H2I4 | SYK | *Taeniopygia guttata* | Chordata | Bird | 2 SH2, 1 Kinase |
| 49 | A0A1V4J5M0 | SYK | *Patagioenas fasciata monilis* | Chordata | Bird | 2 SH2, 1 Kinase |
| 50 | A0A493TXF0 | SYK | *Anas platyrhynchos* | Chordata | Bird | 2 SH2, 1 Kinase |
| 51 | F1N9Y5 | SYK | *Gallus gallus* | Chordata | Bird | 2 SH2, 1 Kinase |
| 52 | A0A218UQF1 | SYK | *Lonchura striata domestica* | Chordata | Bird | 2 SH2, 1 Kinase |
| 53 | XP_026722123.1 | SYK | *Athene cunicularia* | Chordata | Bird | 2 SH2, 1 Kinase |
| 54 | XP_029861296.1 | SYK IsoformX1 | *Aquila chrysaetos chrysaetos* | Chordata | Bird | 2 SH2, 1 Kinase |
| 55 | XP_040396840.1 | SYK IsoformX1 | *Cygnus olor* | Chordata | Bird | 2 SH2, 1 Kinase |
| 56 | KAF1479555.1 | SYK | *Megadyptes antipodes* | Chordata | Bird | 2 SH2, 1 Kinase |
| 57 | U3JQ94 | SYK | *Ficedula albicollis* | Chordata | Bird | 2 SH2, 1 Kinase |
| 58 | A0A1U8BEZ1 | SYK | *Mesocricetus auratus* | Chordata | Mammalia | 2 SH2, 1 Kinase |
| 59 | A0A2K6F665 | SYK | *Propithecus coquereli* | Chordata | Mammalia | 2 SH2, 1 Kinase |
| 60 | P48025 | SYK | *Mus musculus* | Chordata | Mammalia | 2 SH2, 1 Kinase |
| 61 | A0A1S3AT02 | SYK | *Erinaceus europaeus* | Chordata | Mammalia | 2 SH2, 1 Kinase |
| 62 | A0A2I2UZ97 | SYK | *Felis catus* | Chordata | Mammalia | 2 SH2, 1 Kinase |
| 63 | W5PD78 | SYK | *Ovis aries* | Chordata | Mammalia | 2 SH2, 1 Kinase |
| 64 | A0A5F9CUA6 | SYK | *Oryctolagus cuniculus* | Chordata | Mammalia | 2 SH2, 1 Kinase |
| 65 | H0X5F7 | SYK | *Otolemur garnettii* | Chordata | Mammalia | 2 SH2, 1 Kinase |
| 66 | Q64725 | SYK | *Rattus norvegicus* | Chordata | Mammalia | 2 SH2, 1 Kinase |
| 67 | Q00655 | SYK | *Sus scrofa* | Chordata | Mammalia | 2 SH2, 1 Kinase |
| 68 | A0A2K6PUC6 | SYK | *Rhinopithecus roxellana* | Chordata | Mammalia | 2 SH2, 1 Kinase |
| 69 | U3BGC5 | SYK | *Callithrix jacchus* | Chordata | Mammalia | 2 SH2, 1 Kinase |

| | | | | | | |
|---|---|---|---|---|---|---|
| 70 | F7GLV2 | SYK | *Macaca mulatta* | Chordata | Mammalia | 2 SH2, 1 Kinase |
| 71 | A0A2K5KN52 | SYK | *Cercocebus atys* | Chordata | Mammalia | 2 SH2, 1 Kinase |
| 72 | G1MGB0 | SYK | *Ailuropoda melanoleuca* | Chordata | Mammalia | 2 SH2, 1 Kinase |
| 73 | A0A096P223 | SYK | *Papio anubis* | Chordata | Mammalia | 2 SH2, 1 Kinase |
| 74 | XP_023483322.1 | SYK IsoformX1 | *Equus caballus* | Chordata | Mammalia | 2 SH2, 1 Kinase |
| 75 | A0A0D9R4K5 | SYK | *Chlorocebus sabaeus* | Chordata | Mammalia | 2 SH2, 1 Kinase |
| 76 | XP_016816610.1 | SYK | *Pan troglodytes* | Chordata | Mammalia | 2 SH2, 1 Kinase |
| 77 | ERE77472.1 | SYK | *Cricetulus griseus* | Chordata | Mammalia | 2 SH2, 1 Kinase |
| 78 | A0A2K5PKV5 | SYK | *Cebus imitator* | Chordata | Mammalia | 2 SH2, 1 Kinase |
| 79 | A0A6I9L4D8 | SYK | *Peromyscus maniculatus bairdii* | Chordata | Mammalia | 2 SH2, 1 Kinase |
| 80 | G3R4D0 | SYK | *Gorilla gorilla* | Chordata | Mammalia | 2 SH2, 1 Kinase |
| 81 | G3T8M0 | SYK | *Loxodonta africana* | Chordata | Mammalia | 2 SH2, 1 Kinase |
| 82 | P43405 | SYK | *Homo sapiens* | Chordata | Mammalia | 2 SH2, 1 Kinase |

**Table S2:** Accession number of ZAP-70 of different organisms

| Sl. no | Accession number | Gene | Species | Phylum | Class | Domains |
|---|---|---|---|---|---|---|
| 1 | V9KJG4 | ZAP-70 | *Callorhinchus milii* | Chordata | Chondrichthyes | 2 SH2, 1 Kinase |
| 2 | A0A401SFZ6 | ChiPu_0007750 | *Chiloscyllium punctatum* | Chordata | Chondrichthyes | 2 SH2, 1 Kinase |
| 3 | XP_041061522.1 | ZAP-70 | *Carcharodon carcharias* | Chordata | Chondrichthyes | 2 SH2, 1 Kinase |
| 4 | XP_032902714.1 | ZAP-70 | *Amblyraja radiata* | Chordata | Chondrichthyes | 2 SH2, 1 Kinase |
| 5 | XP_038634070.1 | ZAP-70 IsoformX1 | *Scyliorhinus canicula* | Chordata | Chondrichthyes | 2 SH2, 1 Kinase |
| 6 | XP_016896784.1 | ZAP-70 | *Cynoglossus semilaevis* | Chordata | Osteichthyes | 2 SH2, 1 Kinase |
| 7 | A0A672JN05 | LOC115382146 | *Salarias fasciatus* | Chordata | Osteichthyes | 2 SH2, 1 Kinase |
| 8 | XP_005938595.1 | ZAP-70 | *Haplochromis burtoni* | Chordata | Osteichthyes | 2 SH2, 1 Kinase |
| 9 | XP_015221298.1 | ZAP-70 | *Lepisosteus oculatus* | Chordata | Osteichthyes | 2 SH2, 1 Kinase |
| 10 | M3XJU1 | ZAP-70 | *Latimeria chalumnae* | Chordata | Osteichthyes | 2 SH2, 1 Kinase |
| 11 | XP_026179399.1 | ZAP-70 | *Mastacembelus armatus* | Chordata | Osteichthyes | 2 SH2, 1 Kinase |
| 12 | A0A662YJN3 | EOD39_15354 | *Acipenser ruthenus* | Chordata | Osteichthyes | 2 SH2, 1 Kinase |
| 13 | NP_001018425.1 | ZAP-70 | *Danio rerio* | Chordata | Osteichthyes | 2 SH2, 1 Kinase |
| 14 | A0A672LYH7 | LOC107548917 | *Sinocyclocheilus grahami* | Chordata | Osteichthyes | 2 SH2, 1 Kinase |
| 15 | KAG9270088.1 | ZAP-70 | *Astyanax mexicanus* | Chordata | Osteichthyes | 2 SH2, 1 Kinase |
| 16 | XP_026783834.1 | ZAP-70 | *Pangasianodon hypophthalmus* | Chordata | Osteichthyes | 2 SH2, 1 Kinase |

| | | | | | | |
|---|---|---|---|---|---|---|
| 17 | XP_012736872.2 | ZAP-70 | *Fundulus heteroclitus* | Chordata | Osteichthyes | 2 SH2, 1 Kinase |
| 18 | XP_004553860.2 | ZAP-70 | *Maylandia zebra* | Chordata | Osteichthyes | 2 SH2, 1 Kinase |
| 19 | XP_018583813.1 | ZAP-70 | *Scleropages formosus* | Chordata | Osteichthyes | 2 SH2, 1 Kinase |
| 20 | Q6DCV6 | ZAP-70.L | *Xenopus laevis* | Chordata | Amphibia | 2 SH2, 1 Kinase |
| 21 | Q6DF54 | ZAP-70 | *Xenopus tropicalis* | Chordata | Amphibia | 2 SH2, 1 Kinase |
| 22 | A0A6P8S0Q1 | LOC117365392 | *Geotrypetes seraphini* | Chordata | Amphibia | 2 SH2, 1 Kinase |
| 23 | A0A6P7Z494 | LOC115479942 | *Microcaecilia unicolor* | Chordata | Amphibia | 2 SH2, 1 Kinase |
| 24 | XP_040273715.1 | ZAP-70 isoformX2 | *Bufo bufo* | Chordata | Amphibia | 2 SH2, 1 Kinase |
| 25 | XP_029466798.1 | ZAP-70 isoformX1 | *Rhinatrema bivittatum* | Chordata | Amphibia | 2 SH2, 1 Kinase |
| 26 | A0A6J0SJA2 | LOC110072432 | *Pogona vitticeps* | Chordata | Reptile | 2 SH2, 1 Kinase |
| 27 | XP_005281292.1 | ZAP-70 | *Chrysemys picta bellii* | Chordata | Reptile | 2 SH2, 1 Kinase |
| 28 | A0A402FYT7 | ParPi_0025228 | *Paroedura picta* | Chordata | Reptile | 2 SH2, 1 Kinase |
| 29 | XP_024061102.1 | ZAP-70 isoformX1 | *Terrapene carolina triunguis* | Chordata | Reptile | 2 SH2, 1 Kinase |
| 30 | A0A670K6H0 | LOC114588403 | *Podarcis muralis* | Chordata | Reptile | 2 SH2, 1 Kinase |
| 31 | A0A6I9XF86 | ZAP-70 | *Thamnophis sirtalis* | Chordata | Reptile | 2 SH2, 1 Kinase |
| 32 | H0YPY8 | ZAP-70 | *Taeniopygia guttata* | Chordata | Bird | 2 SH2, 1 Kinase |
| 33 | A0A1V4KPZ2 | ZAP-70 | *Patagioenas fasciata monilis* | Chordata | Bird | 2 SH2, 1 Kinase |
| 34 | U3I3S8 | ZAP-70 | *Anas platyrhynchos* | Chordata | Bird | 2 SH2, 1 Kinase |
| 35 | E1BU42 | ZAP-70 | *Gallus gallus* | Chordata | Bird | 2 SH2, 1 Kinase |
| 36 | A0A218UCB5 | ZAP-70 | *Lonchura striata domestica* | Chordata | Bird | 2 SH2, 1 Kinase |
| 37 | XP_026721024.1 | ZAP-70 | *Athene cunicularia* | Chordata | Bird | 2 SH2, 1 Kinase |
| 38 | XP_029889208.1 | ZAP-70 | *Aquila chrysaetos chrysaetos* | Chordata | Bird | 2 SH2, 1 Kinase |
| 39 | XP_040393122.1 | ZAP-70 | *Cygnus olor* | Chordata | Bird | 2 SH2, 1 Kinase |
| 40 | KAF1503043.1 | ZAP-70 | *Megadyptes antipodes* | Chordata | Bird | 2 SH2, 1 Kinase |
| 41 | U3KHA8 | ZAP-70 | *Ficedula albicollis* | Chordata | Bird | 2 SH2, 1 Kinase |
| 42 | A0A3Q0CUC0 | LOC101833852 | *Mesocricetus auratus* | Chordata | Mammalia | 2 SH2, 1 Kinase |
| 43 | A0A2K6GLH5 | ZAP-70 | *Propithecus coquereli* | Chordata | Mammalia | 2 SH2, 1 Kinase |
| 44 | P43404 | ZAP-70 | *Mus musculus* | Chordata | Mammalia | 2 SH2, 1 Kinase |
| 45 | A0A1S3AJ87 | LOC103124953 | *Erinaceus europaeus* | Chordata | Mammalia | 2 SH2, 1 Kinase |
| 46 | M3WD79 | ZAP-70 | *Felis catus* | Chordata | Mammalia | 2 SH2, 1 Kinase |
| 47 | W5PW03 | ZAP-70 | *Ovis aries* | Chordata | Mammalia | 2 SH2, 1 Kinase |
| 48 | G1SPQ8 | ZAP-70 | *Oryctolagus cuniculus* | Chordata | Mammalia | 2 SH2, 1 Kinase |
| 49 | H0XG88 | ZAP-70 | *Otolemur garnettii* | Chordata | Mammalia | 2 SH2, 1 Kinase |
| 50 | A0A0R4J8U1 | ZAP-70 | *Rattus norvegicus* | Chordata | Mammalia | 2 SH2, 1 Kinase |
| 51 | A0A5K1V762 | ZAP-70 | *Sus scrofa* | Chordata | Mammalia | 2 SH2, 1 Kinase |
| 52 | A0A2K6R1D7 | ZAP-70 | *Rhinopithecus roxellana* | Chordata | Mammalia | 2 SH2, 1 Kinase |

| | | | | | | |
|---|---|---|---|---|---|---|
| 53 | F6SWY7 | ZAP-70 | *Callithrix jacchus* | Chordata | Mammalia | 2 SH2, 1 Kinase |
| 54 | F7FAI7 | ZAP-70 | *Macaca mulatta* | Chordata | Mammalia | 2 SH2, 1 Kinase |
| 55 | A0A2K5MYN1 | ZAP-70 | *Cercocebus atys* | Chordata | Mammalia | 2 SH2, 1 Kinase |
| 56 | G1LF11 | ZAP-70 | *Ailuropoda melanoleuca* | Chordata | Mammalia | 2 SH2, 1 Kinase |
| 57 | A0A2I3LEH3 | ZAP-70 | *Papio anubis* | Chordata | Mammalia | 2 SH2, 1 Kinase |
| 58 | F7AR49 | ZAP-70 | *Equus caballus* | Chordata | Mammalia | 2 SH2, 1 Kinase |
| 59 | A0A0D9RXH6 | ZAP-70 | *Chlorocebus sabaeus* | Chordata | Mammalia | 2 SH2, 1 Kinase |
| 60 | H2QIE3 | ZAP-70 | *Pan troglodytes* | Chordata | Mammalia | 2 SH2, 1 Kinase |
| 61 | G3H1Q5 | I79_004087 | *Cricetulus griseus* | Chordata | Mammalia | 2 SH2, 1 Kinase |
| 62 | A0A2K5R1S1 | ZAP-70 | *Cebus imitator* | Chordata | Mammalia | 2 SH2, 1 Kinase |
| 63 | A0A6J0E3Y4 | LOC102923421 | *Peromyscus maniculatus bairdii* | Chordata | Mammalia | 2 SH2, 1 Kinase |
| 64 | G3QGN8 | ZAP-70 | *Gorilla gorilla* | Chordata | Mammalia | 2 SH2, 1 Kinase |
| 65 | G3UMI3 | ZAP-70 | *Loxodonta africana* | Chordata | Mammalia | 2 SH2, 1 Kinase |
| 66 | P43403 | ZAP-70 | *Homo sapiens* | Chordata | Mammalia | 2 SH2, 1 Kinase |

**Table S3**: Receptor conformations used in the model and their corresponding symbols in ODEs

| Conformations | Symbol |
|---|---|
| 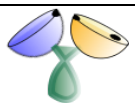 | $R^{00}_{open}$ |
| 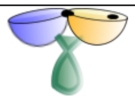 | $R^{00}_{closed}$ |
| 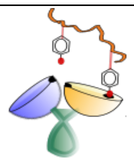 | $R^{01}_{open}$ |
| 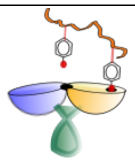 | $R^{01}$ |
| 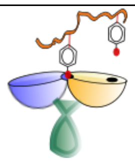 | $R^{10}$ |
| 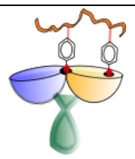 | $R^{11}$ |

**Table S4**: Rate parameters used in the model (Corresponding to Figure 2 b-g)

| **Rate parameters** | **Values** |
|---|---|
| $k_f$ | 0.35 uM$^{-1}$ s$^{-1}$ (varied for Figure 2b,2c) |
| $k_b = k_b'$ | 0.0007 s$^{-1}$ |
| $k_{close}$ | 0.0875 s$^{-1}$ |
| $k_{open}$ | 0.0007 s$^{-1}$ |
| $k_{close}'$ | 8.75×10$^{-8}$ s$^{-1}$ |
| $k_{open}'$ | 0.0014 s$^{-1}$ |
| $k_1^+$ | 0.00035 uM$^{-1}$ s$^{-1}$ |
| $k_1^-$ | 0.00007 s$^{-1}$ |
| $k_2^+$ | 0.00035 uM$^{-1}$ s$^{-1}$ |
| $k_2^-$ | 0.00007 s$^{-1}$ |
| $k_3^+$ | 0.00035 uM$^{-1}$ s$^{-1}$ |
| $k_3^-$ | 0.7×10$^{-9}$ s$^{-1}$ |
| $w_1$ | 0.0002 (varied for Figure 2e, 2f) |
| $w_2$ | 0.02 (Varied for Figure S2a, S2b) |
| $w_3$ | Assumed to be same as w1 |

**Table S5**: Rate parameters and corresponding dissociation constants used in the model to match the experimental data of ITAM-Y2P-ζ1 and ITAM-Y2P-ζ3 bindings to t-SH2 domains of ZAP-70 (corresponding to Figure 2h)

| **Rate parameters** | **ITAM-Y2P-ζ1** | **ITAM-Y2P-ζ3** |
|---|---|---|
| $k_f$ | 1.75 uM$^{-1}$ s$^{-1}$ | 0.35 uM$^{-1}$ s$^{-1}$ |
| $k_b$ | 0.0035 s$^{-1}$ | 0.0007 s$^{-1}$ |
| $k_{close}$ | 0.0875 s$^{-1}$ | 0.00875 s$^{-1}$ |
| $k_{open}$ | 0.0035 s$^{-1}$ | 0.007 s$^{-1}$ |
| $k_{close}'$ | 8.75×10$^{-8}$ s$^{-1}$ | 8.75×10$^{-9}$ s$^{-1}$ |

| | | | |
|---|---|---|---|
| $k'_{open}$ | | 0.00175 $s^{-1}$ | 0.0014 $s^{-1}$ |
| $k_1^+$ | | 0.175 $uM^{-1}$ $s^{-1}$ | 0.035 $uM^{-1}$ $s^{-1}$ |
| $k_1^-$ | | 10.5 $s^{-1}$ | 2.1 $s^{-1}$ |
| $k_2^+$ | | 0.175 $uM^{-1}$ $s^{-1}$ | 0.035 $uM^{-1}$ $s^{-1}$ |
| $k_2^-$ | | 10.5 $s^{-1}$ | 2.1 $s^{-1}$ |
| $k_3^+$ | | 0.00875 $uM^{-1}$ $s^{-1}$ | 0.001225 $uM^{-1}$ $s^{-1}$ |
| $k_3^-$ | | $3.5\times10^{-9}$ $s^{-1}$ | $0.7\times10^{-9}$ $s^{-1}$ |
| $w_1, w_3$ | | $10^{-6}$ | $2\times10^{-7}$ |
| $w_2$ | | $4\times10^{-5}$ | $2\times10^{-5}$ |
| $k'_b$ | | $3.5\times 10^{-6}$ $s^{-1}$ | $7\times10^{-7}$ $s^{-1}$ |
| Dissociation Constants | $K_{d1}(=\frac{k_b}{k_f})$ | 2 nM | 2 nM |
| | $K^*_{d1}$ $(=\frac{k'_b}{w_2 k_f})$ | 50 nM | 100 nM |
| | $K_{d2}$ $(=\frac{k'_b}{w_1 k_f})$ | 2 uM | 10 uM |
| | $K_{con}$ $(=\frac{k_{open}}{k_{close}})$ | 40 nM | 800 nM |
| | $K'_{con}$ $(=\frac{k'_{open}}{k'_{close}})$ | 20000 | 20000 |

**Table S6**: Rate parameters used in the model (shown in Figure 3f) to match the experimental data of tSH2: ITAM-YP-ζ1 and $^{tSH2}$R39A: ITAM-Y2P-ζ1 (Figure 3 g-h)

| Rate parameters | tSH2: ITAM-YP-ζ1 | $^{tSH2}$R39A: ITAM-Y2P-ζ1 |
|---|---|---|
| $k_f$ | 0.875 $uM^{-1}$ $s^{-1}$ | 0.4375 $uM^{-1}$ $s^{-1}$ |
| $k_b$ | 0.0035 $s^{-1}$ | 0.0035 $s^{-1}$ |
| $k_{close}$ | 0.875 $s^{-1}$ | 0.875 $s^{-1}$ |
| $k_{open}$ | 0.0875 $s^{-1}$ | 2.1875 $s^{-1}$ |

| | | | |
|---|---|---|---|
| $w_1, w_2, w_3, k_b'$ , $k_1^+, k_1^-, k_2^+, k_2^-, k_3^+, k_3^-$, $k_{close}'$ , $k_{open}'$ (negligible) | | $10^{-10}$ unit | $10^{-10}$ unit |
| Dissociation Constants | $K_{d1}(=\frac{k_b}{k_f})$ | 4 nM | 8 nM |
| | $K_{con}$ $(=\frac{k_{open}}{k_{close}})$ | 100 nM | 2.5 uM |

**Table S7**: Rate parameters used in the model (shown in Figure 6h) to match the experimental data of Syk tSH2 binding (Figure 6i-j). Note, the parameters corresponding to ITAM-YP-ζ1 and ITAM-YP-ζ3 are the same since Syk cannot possibly distinguish between them according to the experiments (Figure 6a, 6i). We only varied $K_d = \frac{k_b}{k_f}$ by changing $k_b$, and $K_d$ = 60 nM produced the best fit to the data.

| Rate symbols | Values (both for ITAM-Y2P-ζ1 and ITAM-Y2P-ζ3) |
|---|---|
| $k_f$ | 1.3 $uM^{-1}$ $s^{-1}$ |
| $k_{close}$ | 1.3 $s^{-1}$ |
| $k_b$ | 0.026 $s^{-1}$, 0.078 $s^{-1}$,0.13 $s^{-1}$ ($K_d = \frac{k_b}{k_f}$ = 20 nM,60 nM,100 nM) |
| $k_{open}$ (=$k_b$) | Assumed to be the same as $k_b$ |
| $w_1, w_2, w_3$ | 1 (no kinetic penalty) |
| $k_1^+, k_1^-, k_2^+, k_2^-, k_3^+, k_3^-$, $k_{close}'$ , $k_{open}'$ (negligible) | $10^{-10}$ unit |

#### References:


1 Gangopadhyay, K., Manna, B., Roy, S., Kumari, S., Debnath, O., Chowdhury, S., Ghosh, A. and Das, R. (2020) An allosteric hot spot in the tandem-SH2 domain of ZAP-70 regulates T-cell signaling. Biochemical Journal. **477**, 1287-1308

2 Mayer, B. J., Jackson, P. K. and Baltimore, D. (1991) The noncatalytic src homology region 2 segment of abl tyrosine kinase binds to tyrosine-phosphorylated cellular proteins with high affinity. Proceedings of the National Academy of Sciences of the United States of America. **88**, 627-631

3 Machida, K., Thompson, C. M., Dierck, K., Jablonowski, K., Karkkainen, S., Liu, B., Zhang, H., Nash, P. D., Newman, D. K., Nollau, P., Pawson, T., Renkema, G. H., Saksela, K., Schiller, M. R., Shin, D. G. and Mayer, B. J. (2007) High-throughput phosphotyrosine profiling using SH2 domains. Molecular cell. **26**, 899-915

4 Suzuki, Y., Moriyoshi, E., Tsuchiya, D. and Jingami, H. (2004) Negative cooperativity of glutamate binding in the dimeric metabotropic glutamate receptor subtype 1. The Journal of biological chemistry. **279**, 35526-35534

5 Chakraborty, M. P., Bhattacharyya, S., Roy, S., Bhattacharya, I., Das, R. and Mukherjee, A. (2021) Selective targeting of the inactive state of hematopoietic cell kinase (Hck) with a stable curcumin derivative. The Journal of biological chemistry. **296**, 100449

6 Consortium, T. U. (2020) UniProt: the universal protein knowledgebase in 2021. Nucleic Acids Research. **49**, D480-D489

7 Altschul, S. F., Madden, T. L., Schäffer, A. A., Zhang, J., Zhang, Z., Miller, W. and Lipman, D. J. (1997) Gapped BLAST and PSI-BLAST: a new generation of protein database search programs. Nucleic Acids Research. **25**, 3389-3402

8 (2018) Database resources of the National Center for Biotechnology Information. Nucleic Acids Res. **46**, D8-d13

9 de Castro, E., Sigrist, C. J., Gattiker, A., Bulliard, V., Langendijk-Genevaux, P. S., Gasteiger, E., Bairoch, A. and Hulo, N. (2006) ScanProsite: detection of PROSITE signature matches and ProRule-associated functional and structural residues in proteins. Nucleic Acids Res. **34**, W362-365

10 Felsenstein, J. (1985) CONFIDENCE LIMITS ON PHYLOGENIES: AN APPROACH USING THE BOOTSTRAP. Evolution. **39**, 783-791

11 Jones, D. T., Taylor, W. R. and Thornton, J. M. (1992) The rapid generation of mutation data matrices from protein sequences. Comput Appl Biosci. **8**, 275-282

12 Kumar, S., Stecher, G., Li, M., Knyaz, C. and Tamura, K. (2018) MEGA X: Molecular Evolutionary Genetics Analysis across Computing Platforms. Molecular biology and evolution. **35**, 1547-1549

13 Le, S. Q. and Gascuel, O. (2008) An Improved General Amino Acid Replacement Matrix. Molecular Biology and Evolution. **25**, 1307-1320

14 Letunic, I., Khedkar, S. and Bork, P. (2020) SMART: recent updates, new developments and status in 2020. Nucleic Acids Research. **49**, D458-D460

15 Crooks, G. E., Hon, G., Chandonia, J. M. and Brenner, S. E. (2004) WebLogo: a sequence logo generator. Genome research. **14**, 1188-1190

16 Schneider, T. D. and Stephens, R. M. (1990) Sequence logos: a new way to display consensus sequences. Nucleic Acids Research. **18**, 6097-6100

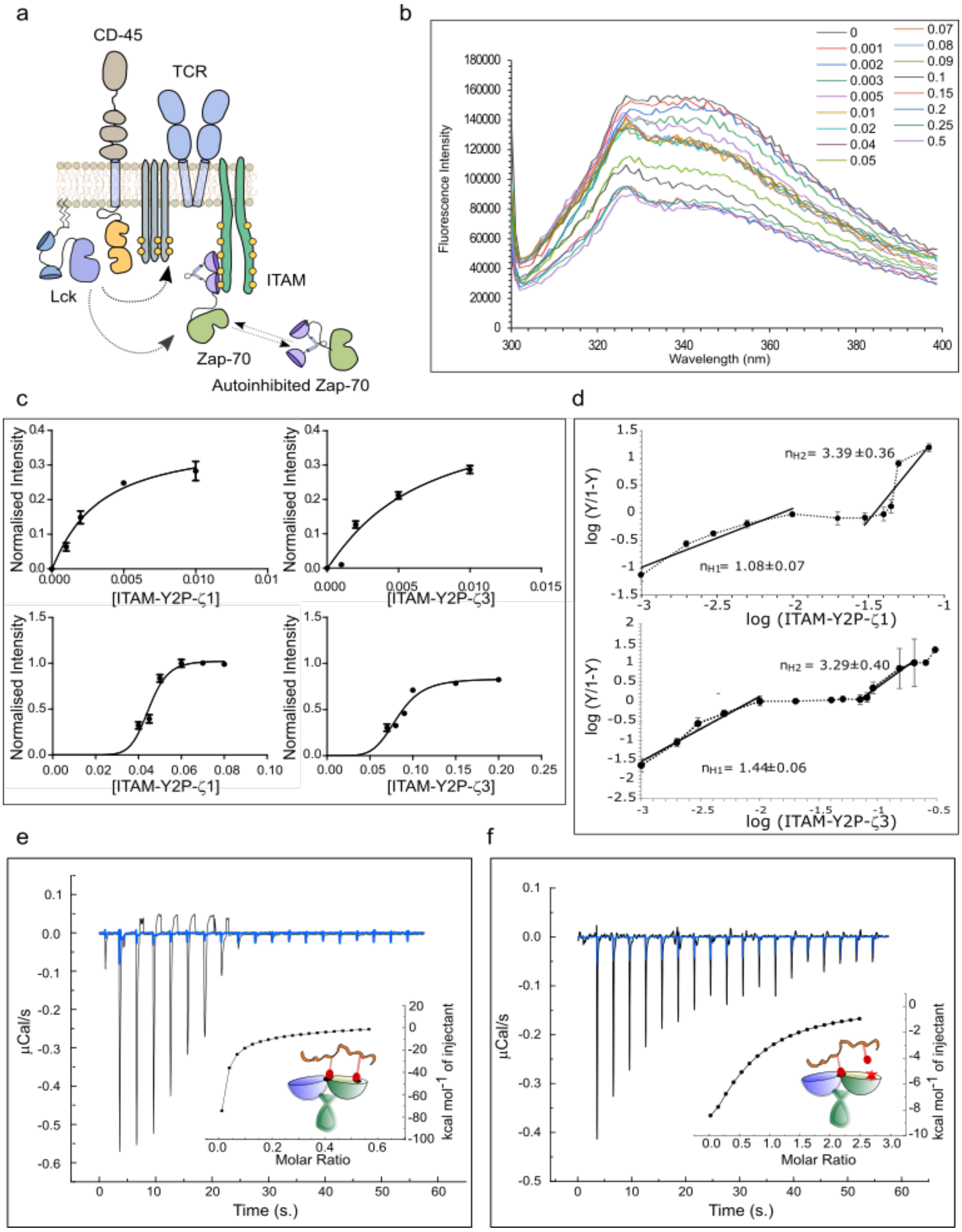


**Figure S1 Binding of tSH2 domain of ZAP-70 to different ITAM motifs (related to Figure 1):** (a) Schematic diagram showing the initiation of T-cell signaling. The Lck Kinase phosphorylates the repeated ITAM motif in the cytosolic domain of T-cell receptor. tSH2 domain of ZAP-70 binds to the doubly phosphorylated ITAM motifs, thus initiates the downstream signaling. (b) Representative fluorescence intensity scanned (λem 300nm to 400nm) from the titration of tSH2 domain of ZAP-70 and ITAM-Y2P-ζ3 peptide. (c) The change in intrinsic fluorescence for tSH2 domain of ZAP-70 with increasing concentration of

ITAM-Y2P-ζ1 and ITAM-Y2P-ζ3, respectively (Figure 1d) was fitted to first-order binding equation implemented in the program Prism. (d) Hill plot for the titration of tSH2 domain of ZAP-70 and different peptides (ITAM-Y2P-ζ1 and ITAM-Y2P-ζ3), described in figure 1d. The Hill-coefficient (nH) was determined from the slope of the plot. (e-f) Representative isothermal titration calorimetry for the tSH2 domain and ITAM-Y2P-ζ3 peptide, and $^{tSH2}$R190A and ITAM-Y2P-ζ3 peptide, respectively.

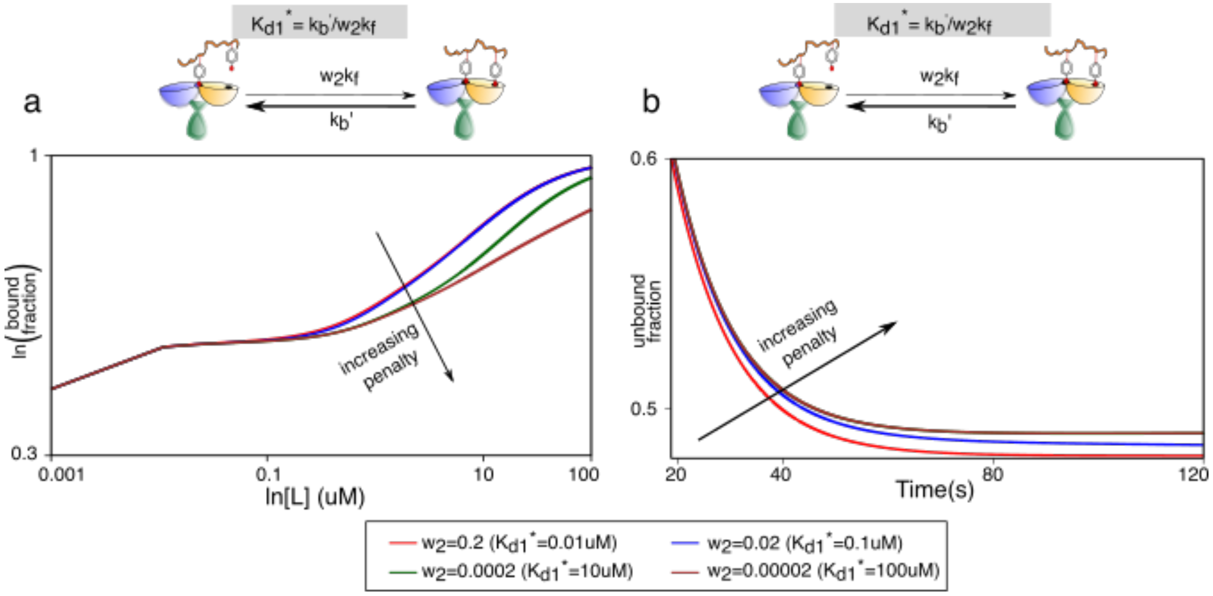


**Figure S2 (Related to Figure 2): Variation of $K_{d1}^{*}$ does not significantly affect the plateau width and the kinetic profiles in the steady state and pre-steady state conditions, respectively.** (a) Effects of $K_{d1}^{*}$ variation (by changing the penalty factor $w_2$) on the bound fraction under steady state. (b) Effects of $K_{d1}^{*}$ variation on kinetics of the unbound fraction showing only a single-step decay.

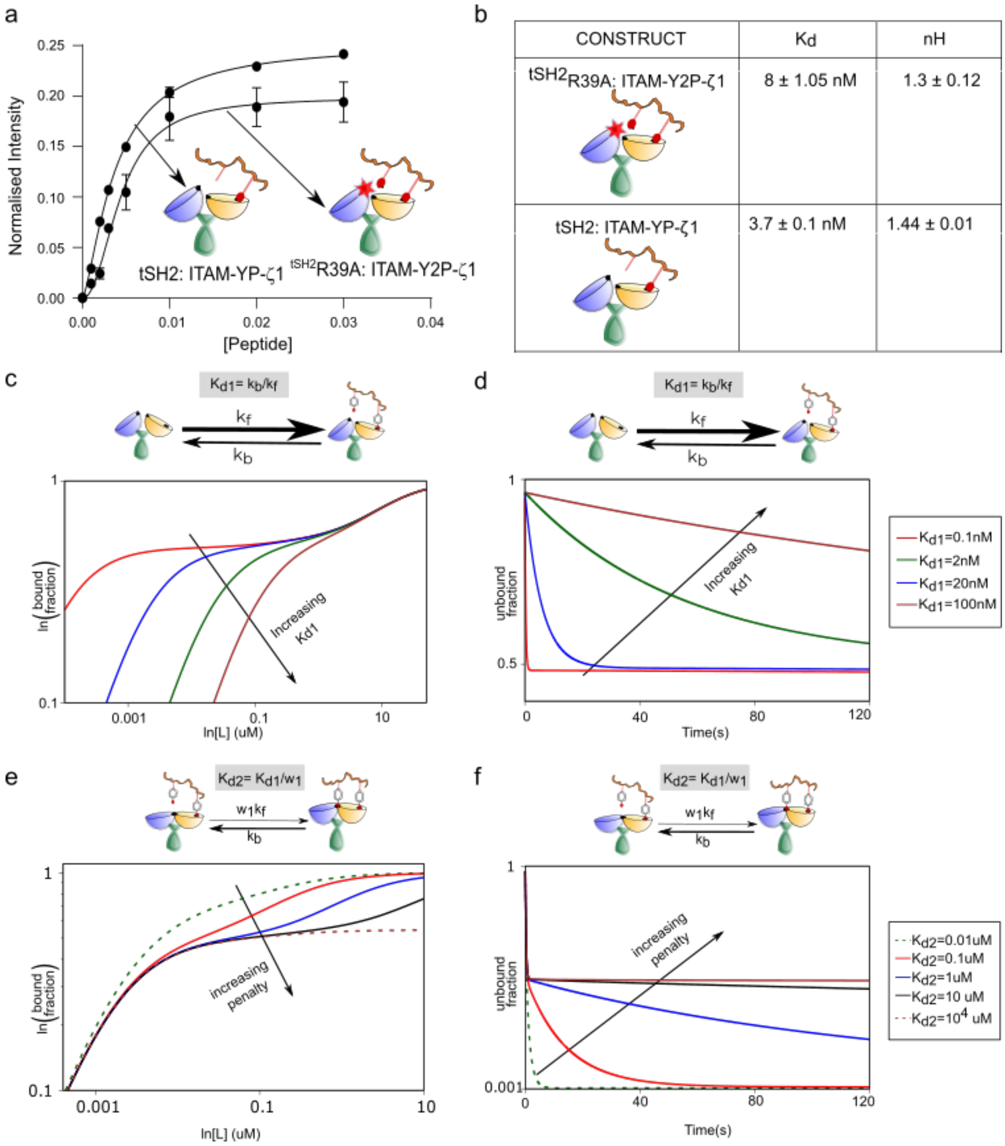


**Figure S3 (Related to Figure 3):** (a) The change in intrinsic fluorescence for $^{tSH2}$R39A: ITAM-Y2P-ζ1 and tSH2 wildtype: ITAM-YP-ζ1 titration, respectively, was fitted to first-order binding equation implemented in the program Prism. (b) Tabulation of the $K_d$ and Hill coefficient (nH) of the binding of $^{tSH2}$R39A and ITAM-Y2P-ζ1, and tSH2 wildtype and ITAM-YP-ζ1, respectively. (c-d) Effects of $K_{d1}$ variation in the reduced model shown in Figure 3k. Similar to the results of the full model (Figure 2b-g), the $K_{d1}$ determines the sensitivity of the bound-fraction in the steady-state (panel c) and modulates the initial decay in the kinetics of unbound fraction (panel d). (e) Increasing $K_{d2}$ broadens the plateau width in the steady state, similar to the full model. (f) Binding kinetics again show a two-step decay as in the full model.

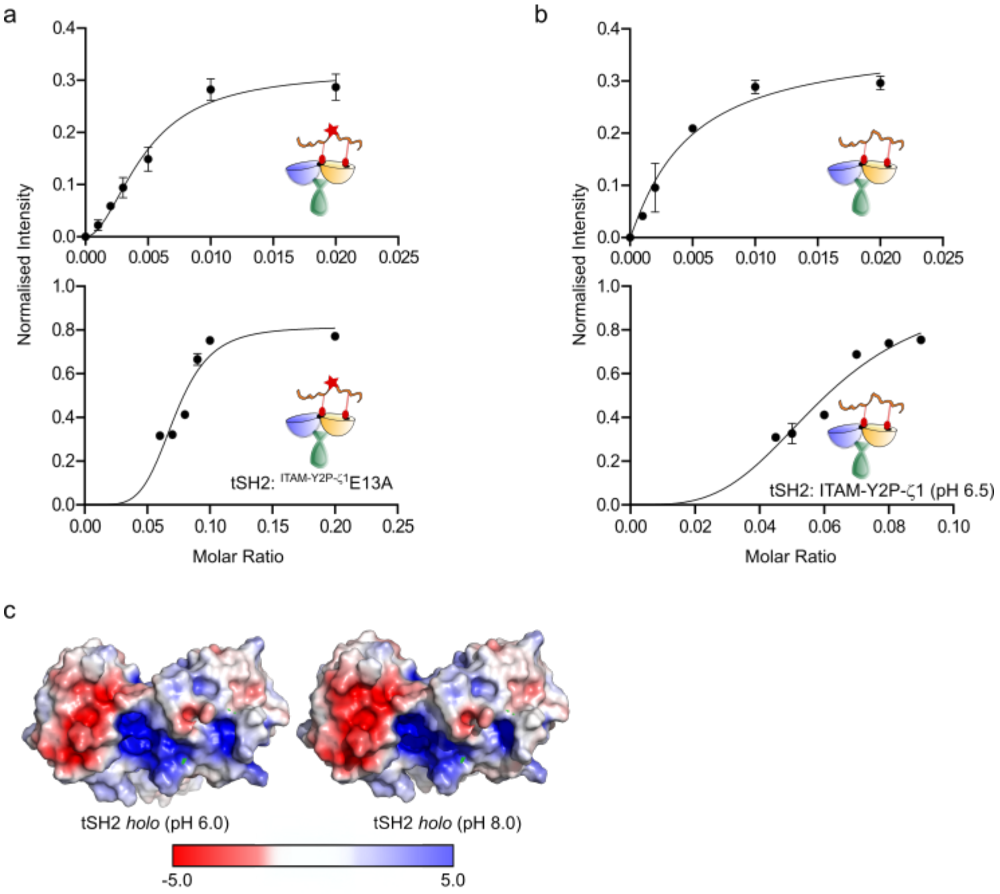


**Figure S4 (Related to Figure 4):** (a-b) The change in intrinsic fluorescence for tSH2 domain of ZAP-70: $^{\text{ITAM-Y2P-}\zeta 1}$E13A at pH 8.0, and ITAM-Y2P-ζ1 at pH 6.5, respectively, was fitted to first-order binding equation implemented in the program Prism. (c) Surface potential for the tSH2 *holo* structure of ZAP-70 (PDB ID: 2OQ1) at pH 6.0 and pH 8.0, respectively, are shown. Color bar represents charge density. (d) $\Delta\Delta G_{\text{unfolding}}$ calculation at 40˚C for tSH2 domain of ZAP-70 bound with ITAM-Y2P-ζ1, ITAM-Y2P-ζ3, and $^{\text{ITAM-Y2P-}\zeta 1}$E13A. The calculations were done in respect to the $\Delta G_{\text{unfolding}}$ value of tSH2 *apo* state at 40˚C.

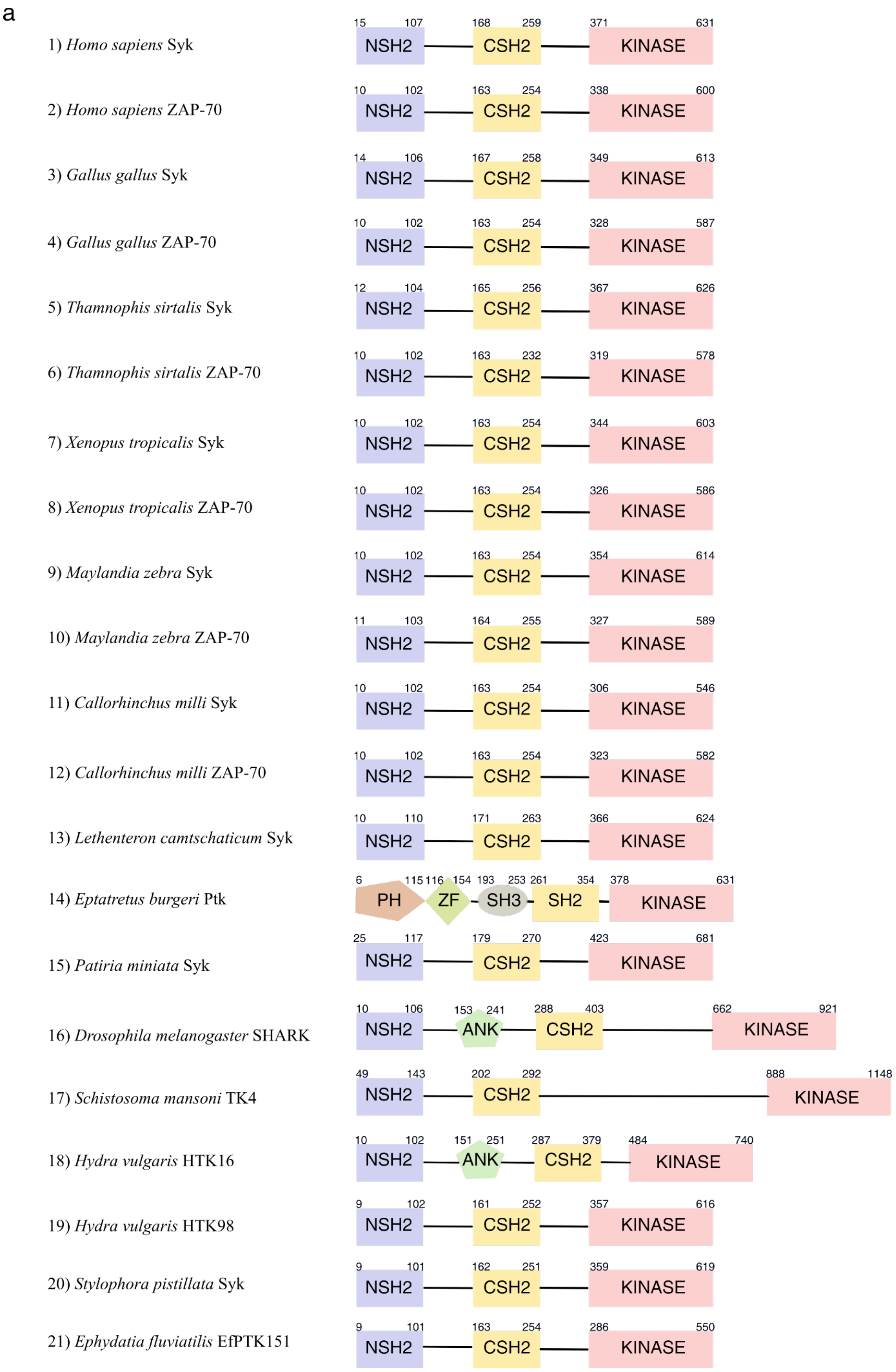


**Figure S5: Domain Architecture of Syk Family Kinases (Related to Figure 5)**

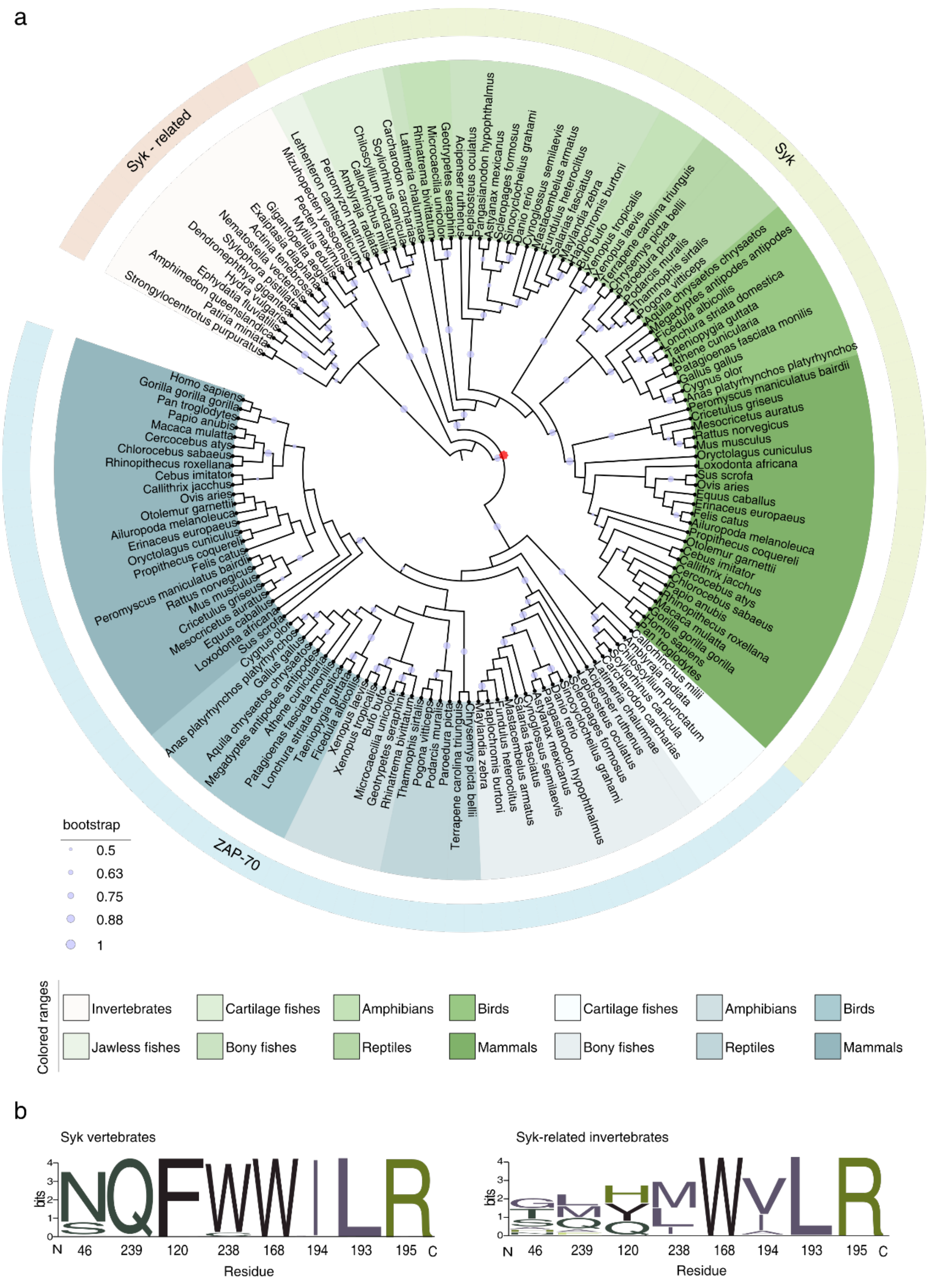


**Figure S6: (Related to Figure 5):** (a) The phylogenetic tree represents the evolutionary relationship between Syk, Syk-related kinases, and ZAP-70. The color code represents the classes to which each species belongs. The red dot depicts the emergence of ZAP-70 in jawed vertebrates from cartilage fish to mammals. (b) Sequence logo showing the conservation of network residues of Syk in vertebrates(left) and invertebrates(right). Except for N46, and W238 all other residues are conserved in vertebrates. The Syk-related kinases show high residue variation in invertebrates.

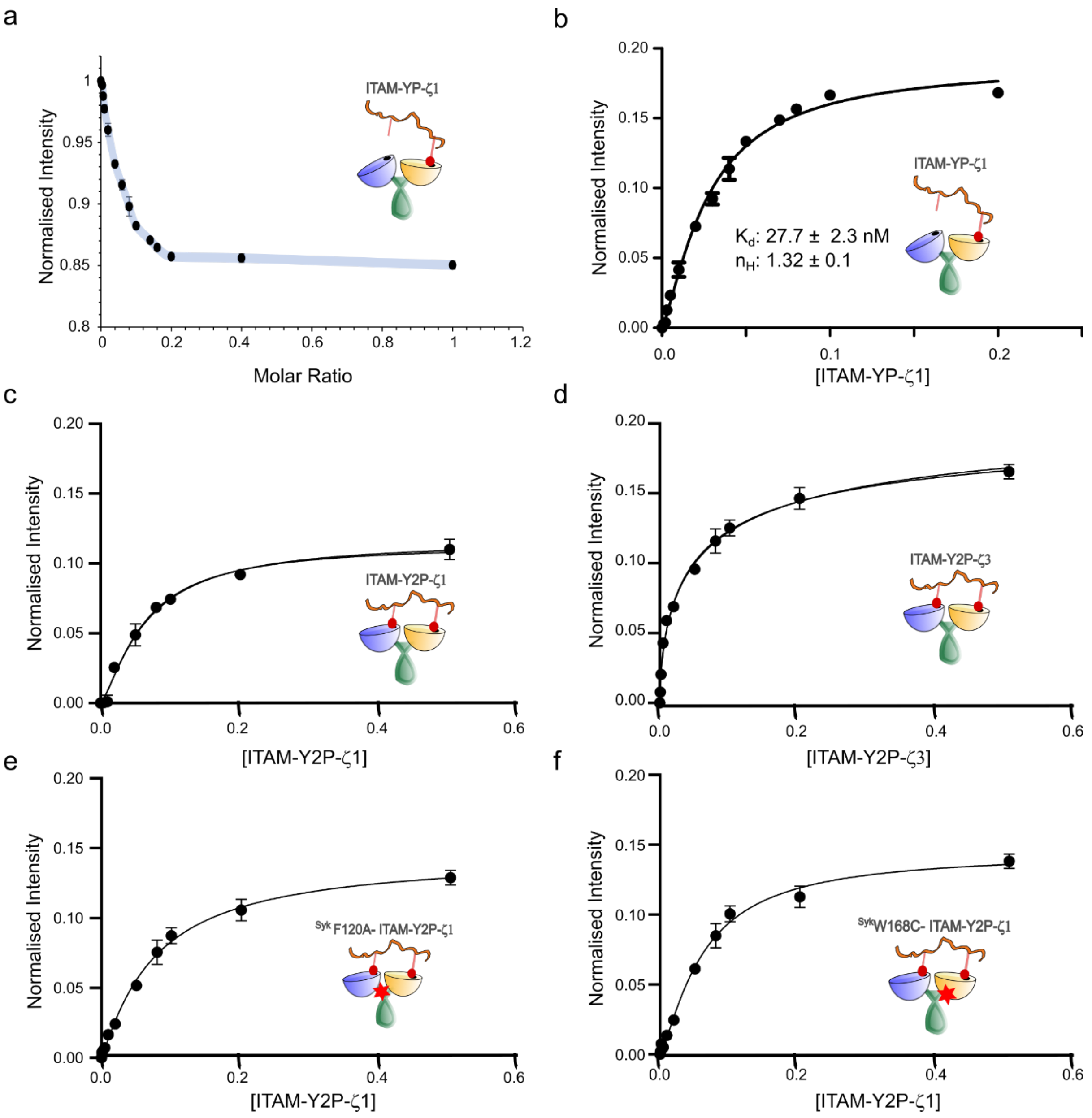


**Figure S7: Binding of Syk tSH2 domain and mutants and doubly-phosphorylated ITAM-Y2P peptides (Related to Figure 6):** (a-b) Titration of ITAM-YP-ζ1 and tSH2 domain of Syk determined from the measurement of intrinsic tryptophan-fluorescence at the indicated ligand to protein molar ratio. The error bar represents the standard deviation from three experiments. (c- d) The change in intrinsic fluorescence for tSH2 domain of Syk with increasing concentration of ITAM-Y2P-ζ1and ITAM-Y2P-ζ3 (Figure 5a) was fitted to first-order binding equation implemented in the program Prism. (e-f) Syk tSH2 mutants (F120A and W168C) was titrated with increasing concentration of ITAM-Y2P-ζ1 and the data was fitted to first-order binding equation using Prism.
